# Structural architecture and brain network efficiency links polygenic scores to intelligence

**DOI:** 10.1101/2022.03.22.485284

**Authors:** Erhan Genç, Dorothea Metzen, Christoph Fraenz, Caroline Schlüter, Manuel C. Voelkle, Larissa Arning, Fabian Streit, Huu Phuc Nguyen, Onur Güntürkün, Sebastian Ocklenburg, Robert Kumsta

**Author notes:** These authors contributed equally (Erhan Genç, Dorothea Metzen). These authors are joint senior authors (Sebastian Ocklenburg, Robert Kumsta).

## Abstract

Intelligence is highly heritable. Genome-wide association studies (GWAS) have shown that thousands of alleles contribute to variation in intelligence with small effect sizes. Polygenic scores (PGS), which combine these effects into one genetic summary measure, are increasingly used to investigate polygenic effects in independent samples. Whereas PGS explain a considerable amount of variance in intelligence, it is largely unknown how brain structure and function mediate this relationship. Here we show that individuals with higher PGS for educational attainment and intelligence had higher scores on cognitive tests, larger surface area, and more efficient fiber connectivity derived by graph theory. Fiber network efficiency as well as surface of brain areas partly located in parieto-frontal regions were found to mediate the relationship between PGS and cognitive performance. These findings are a crucial step forward in decoding the neurogenetic underpinnings of intelligence, as they identify specific regional networks that link polygenic predisposition to intelligence.

## Introduction

Intelligence is a general mental capability that involves the ability to reason, plan, solve problems, and learn from experience (*1*). General intelligence, or *g*, is one of the most intensely studied psychological phenotypes for its high stability across the life course (*2*) and its high predictive value for educational success (*3*) and health outcomes (*4*). Despite intelligence’s high relevance in everyday life, investigating its neurogenetic underpinnings showed to be surprisingly challenging (*5*).

Intelligence is a highly heritable trait (*5*), with about 50% of variance accounted for by genetic factors. Genome wide association studies (GWAS), which test the association between single nucleotide polymorphisms (SNPs) with a phenotype, showed that intelligence is highly polygenic, with thousands of alleles across the genome contributing with small effect sizes (*6*). One way forward in accounting for this highly polygenic architecture is to combine effects of different SNPs across the whole genome into one summary measure, so-called polygenic scores (PGS) (*7*). PGS are determined by computing the sum of allelic effects for a specific phenotype such as intelligence over the whole genome and weighting them with an effect size estimate obtained from GWAS. Importantly, PGS use the statistical power of well-powered GWAS of discovery samples to be applied robustly in smaller target samples (*8, 9*). In the case of intelligence, PGS derived from one of the largest GWAS to date (*6*) explain up to 5.2% of variance in general intelligence. For educational attainment - highly correlated to intelligence and much easier to measure - larger GWAS could be realized, with resulting PGS that explain up to 11% of variance in educational attainment (*10*), and 7% of IQ variance (*5*).

In addition to their potential diagnostic utility, PGS can be leveraged to map the pathway from genetic disposition to phenotype. Whereas it is known that intelligence is influenced by brain structure and function as well as network efficiency (*11, 12*), a functional understanding of which specific brain parameters mediate the link between genetic variation and intelligence is missing. Several brain properties are related to intelligence, including brain volume, surface area, and cortical thickness (*13–16*). Importantly, intelligence is not tied to the properties of one single brain area, but to a wide network of brain areas spread across the whole cortex. Here, a network mainly comprising the dorsolateral prefrontal cortex, the parietal lobe, the anterior cingulate cortex, the temporal lobe, and the occipital lobe seems to be central for cognitive performance, as proposed by the Parieto-Frontal Integration Theory of intelligence (P-FIT) (*17*). The theory assumes that all of these P-FIT areas, even though they were identified independently of each other, are likely to have strong interconnections and form an extensive brain network. Recent studies and models focusing on connectivity-based approaches indicate that there may be brain areas whose structural and functional properties are not related to intelligence, while their connectivity patterns are (*12, 18*). Previous research, in which the connectivity between brain regions was quantified via diffusion-weighted imaging (DWI) and graph theoretical approaches, showed that the brain’s global efficiency as well as the nodal efficiency of brain areas from the P-FIT network and beyond are associated with intelligence (*19–25*). In addition to structural connectivity, graph theory can also be used in combination with data from resting-state fMRI in order to study the brain’s functional connectivity (*26*). There is evidence that general intelligence is positively correlated with functional global efficiency (*27*) and the nodal efficiency of areas belonging to the P-FIT network. However, subsequent studies could not replicate these associations (*28–30*).

Thus, macrostructural properties of specific brain areas and the structural efficiency of the human connectome represent likely candidates for mediating the effects of genetic variation on general intelligence. Several GWAS reporting genetic correlations between brain properties and intelligence, i.e. overlapping genetic variants being associated with both phenotypes, support this notion (*31–37*). In a complementary approach, studies demonstrated associations between PGS for educational attainment or general intelligence and brain properties (*38–40*). However, true mediation analyses are rare. A true mediation analysis requires the polygenic disposition, brain properties (putative mediator) and intelligence (outcome) to be measured in the same sample. By doing so, one can directly analyze the extent to which the association between PGS and intelligence is explained via variation in brain structure and function. Three studies to date have investigated the mediation effect on the macrostructural level (*41–43*). Elliot *et al*. (*41*) analyzed potential mediation effects of total brain volume on the relationship between PGS for educational attainment and cognitive performance. They found that participants with larger brains and with higher PGS performed better on cognitive tests. PGS were also positively associated with brain size. However, there was no clear overall mediation effect of brain volume. Since general intelligence is associated with specific regions in the brain, subsequent studies focused on region-specific mediation effects of cortical thickness and surface area. Lett *et al.* (*42*) employed PGS for general intelligence and found that the association between PGS and general intelligence was partially mediated by surface area and cortical thickness in prefrontal regions, anterior cingulate, insula, and medial temporal cortex. It is noteworthy that some of these regions are part of the P-FIT network. Results were consistent across two independent samples, indicating that macrostructural properties of specific areas, partly belonging to the P-FIT network, may indeed play a crucial role with regard to the link between genetic variation and general intelligence. Another study by Mitchell *et al.* (*43*), which employed PGS for educational attainment, reported similar findings. They observed that surface area and cortical thickness of specific cortical regions partially mediated the effects of PGS on cognitive test performance. These regions were the fusiform gyrus, entorhinal cortex, banks of the superior temporal sulcus, the inferior frontal gyrus, and the medial orbital frontal gyrus.

To summarize, there is evidence that specific gray matter macrostructural properties of brain areas from the P-FIT network represent likely candidates to explain the link between genetic variation and intelligence. What is missing, however, is a systems-view taking into account white matter connectivity as well as functional network properties. Our study aimed to fill this crucial gap in the literature by using a multilevel deep phenotyping approach, including an integrated analysis of behavioral and neuroimaging phenotypes. We investigated the effects of two different PGS on general intelligence: PGS for educational attainment (*10*) and PGS for general intelligence (*6*). We tested the mediating role of surface area, cortical thickness, white matter fiber network efficiency, and functional network efficiency on the level of the whole brain as well as for specific brain areas. Thus, this study presents the first multimodal mediation analysis that gives brain region-specific insight into the putative links between genetics and general intelligence.

## Methods

### Participants

Our sample consisted of 557 adults, who reported to be free from past or present neurological and/or psychological conditions. The mean age was 27.33 years (SD = 9.43; range = 18-75), we tested 283 men (mean age = 27.1, SD = 9.86) and 274 women (mean age = 26.94, SD = 8.96). Participants were mostly university students (mean years of education = 17.4, SD = 3.12), who participated in exchange for course credit or financial compensation. The study was approved by the local ethics committee of the Faculty of Psychology at Ruhr-University Bochum (Nr. 165). All participants gave written informed consent and were treated according to the Declaration of Helsinki. The final dataset (see Statistical analysis) included 523 participants aged from 18 to 75 (M = 27.1, SD = 9.08, 266 women). The data is part of a large-sample study on the neural correlates of intelligence, personality, and motivation. Hence, it has been used in other publications (*44–46*).

### General intelligence testing: I-S-T 2000 R

Since participants were native German speakers, general intelligence was assessed using the “Intelligenz-Struktur-Test 2000 R” (I-S-T 2000 R), a well-established German intelligence test battery (*47, 48*). The test was conducted in a quiet and well-lit room. The I-S-T 2000 R comprises various types of mental test items to measure multiple facets of general intelligence and is largely comparable to the internationally established Wechsler Adult Intelligence Scale (WAIS IV) (*49*). It consists of a basic and an extension module. In this study, participants only completed the basic module. The basic module contains 180 items assessing three sub-factettes of general intelligence, namely verbal, numeric, and figural reasoning (*46*). For every participant a sum score across all items was computed and used as outcome in the mediation analysis.

### Genotyping and polygenic scores (PGS)

Exfoliated cells brushed from the oral mucosa were used for genotyping. DNA isolation was conducted with QIAamp DNA mini Kit (Qiagen GmbH, Hilden, Germany). Genotyping was performed with the Illumina Infinium Global Screening Array 1.0 with MDD and Psych content (Illumina, San Diego, CA, USA) at the Life & Brain facilities (Bonn, Germany). Filtering was done with PLINK 1.9 by eliminating all SNPs with a minor allele frequency of < 0.01, missing data > 0.02, or deviating from Hardy-Weinberg equilibrium by a *p* value < 1*10^-6^. Subjects were excluded due to sex-mismatch, > 0.02 missingness, and heterozygosity rate > |0.2|. A high quality (HWE *p* > 0.02, MAF > 0.02, missingness = 0) and LD pruned (*r*^*2*^ = 0.01) SNP set was used for assessing relatedness and population structure. Pi hat > 0.2 was used to exclude subjects randomly in pairs of related subjects. Finally, we computed principal components to control for population stratification. Individuals who deviated more than 6 SD from the first 20 PCs were categorized as outliers and excluded. The final data set consisted of 523 participants and 492,348 SNPs.

We calculated genome-wide polygenic scores (PGS) for all participants using two publicly available summary statistics: general intelligence (GI, N = 269,867) (*6*) and educational attainment (EA, N = 1,131,881) (*10*). PGS were calculated as weighted sums of a subject’s trait-associated alleles across all SNPs using PRSice 2.1.6. We report the best-fit PGS, meaning that the *p* value threshold for PGS calculation was chosen empirically (in steps of 5*10^-5^ from 5*10^-8^ to 0.5) so that the calculated PGS explained the maximum amount of I-S-T 2000 R variance (*46*). The best-fit threshold selected for PGS_EA_ was 1, for PGS_GI_ it was 0.0062. The statistic “incremental R^2^” was taken as a value for the predictive power of the PGS. Incremental R^2^ stands for the increase in determination coefficient R^2^ when the corresponding PGS is added to a regression model predicting I-S-T 2000 R together with our control variables. The control variables chosen were age, sex, and the first four principal components of population stratification. We used linear parametric methods for all statistical analysis in PRSice. Testing was two-tailed (a-level of *p* < 0.05). PGS_EA_ explained 3.3% of variance in I-S-T 2000 R score, PGS_GI_ explained 4.8%.

### Neuroimaging

#### Acquisition of anatomical data

Magnetic resonance imaging was performed on a 3T Philips Achieva scanner with a 32-channel head coil. The scanner was located at Bergmannsheil University Hospital in Bochum, Germany. T1-weighted data were obtained by means of a high-resolution anatomical imaging sequence with the following parameters: MP-RAGE; TR = 8.179 ms; TE = 3.7 ms; flip angle = 8°; 220 slices; matrix size = 240 × 240; resolution = 1 mm × 1 mm × 1 mm; acquisition time = 6 minutes.

#### Acquisition of diffusion-weighted data

Diffusion-weighted images (DWI) were acquired using echo planar imaging with the following parameters: TR = 7652 ms, TE = 87 ms, flip angle = 90°, 60 slices, matrix size = 112 × 112, resolution = 2 mm × 2 mm × 2 mm. Diffusion weighting was carried out along 60 isotropically distributed directions with a b-value of 1000 s/mm². In addition, six volumes with a b-value of 0 s/mm² and no diffusion weighting were acquired. These served as an anatomical reference for motion correction. In total, we acquired three consecutive scans, which were averaged following established protocol (*44*). This was done to increase the signal-to-noise ratio. Acquisition time was 30 minutes.

#### Acquisition of resting-state data

Functional MRI resting-state images were acquired using echo planar imaging (TR = 2000 ms, TE = 30 ms, flip angle = 90°, 37 slices, matrix size = 80 × 80, resolution = 3 mm × 3 mm × 3 mm). Participants were instructed to lay still with their eyes closed and to think of nothing in particular. Acquisition time was 7 minutes.

### Analysis of imaging data

#### Analysis of anatomical data

Cortical surfaces of T1-weighted images were reconstructed using FreeSurfer (http://surfer.nmr.mgh.harvard.edu, version 5.3.0), following established protocol (*50, 51*). Pre-processing included skull stripping, gray and white matter segmentation as well as reconstruction and inflation of the cortical surface. These steps were performed individually for each participant. Slice by slice quality control was performed and inaccuracies of automatic pre-processing were edited manually. For the purpose of brain segmentation, we used the Human Connectome Project’s multi-modal parcellation (HCPMMP). Respective parcellation comprises 180 areas per hemisphere and is based on structural, functional, topographical, and connectivity data of healthy participants (*52*). The original data provided by the Human Connectome Project were converted to annotation files matching the standard cortical surface in FreeSurfer called fsaverage. This fsaverage parcellation was transformed to each participant’s individual cortical surface and converted to volumetric masks. Supplementary to the HCPMMP, eight subcortical gray matter structures per hemisphere were added to the parcellation (thalamus, caudate nucleus, putamen, pallidum, hippocampus, amygdala, accumbens area, ventral diencephalon) (*53*). Finally, six regions representing the four ventricles of the brain were delineated to serve as a reference for later BOLD signal analyses. All masks were linearly transformed into the native spaces of the resting-state and diffusion-weighted images und used as landmarks for graph theoretical connectivity analyses (see Figure 1). As we did not have any specific hypotheses with regard to hemispheric differences, we computed a mean value for each brain region by averaging values across the left and right hemispheres (e.g., the value for area V1 is the mean of L_V1 and R_V1). This resulted in 180 cortical and 8 subcortical areas.

**Figure 1.**
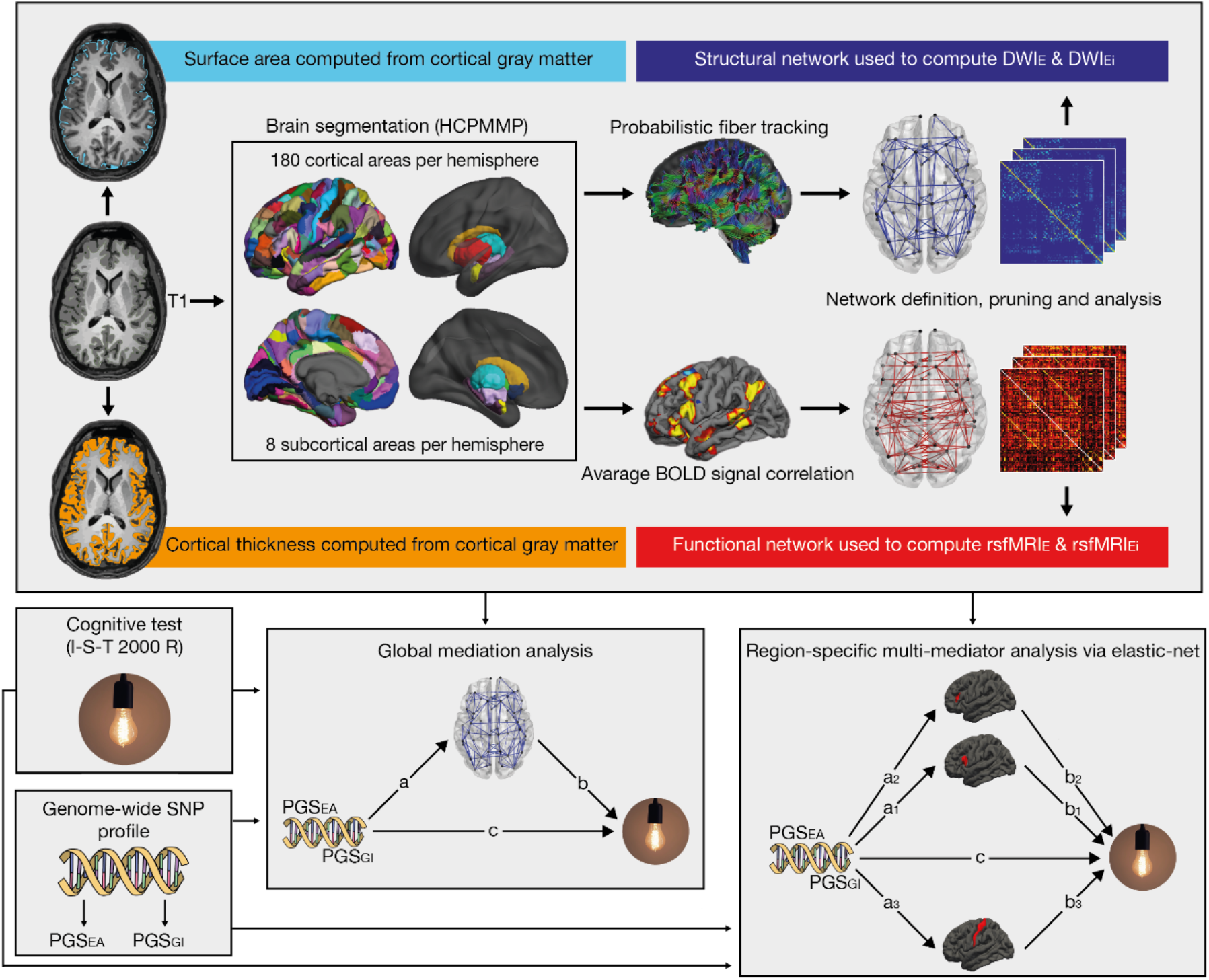
Processing steps of neurocognitive data and statistical analysis. First, T1-weighted anatomical images were used to compute estimates of cortical surface area and cortical thickness. Second, T1-weighted anatomical images were segmented into 180 cortical structures per hemisphere according to the HCPMMP atlas and 8 subcortical structures per hemisphere. Third, the resulting masks were linearly transformed into the native spaces of the resting-state and diffusion-weighted images. For the diffusion-weighted images, probabilistic fiber tracking was carried out with aforementioned masks serving as seed and target regions. For the resting-state images, partial correlations between average BOLD time courses of all brain regions were computed. Fourth, structural and functional networks were constructed. Edges were weighted by the results of probabilistic fiber tractography or BOLD signal correlation. Fifth, these networks were used for the computation of global efficiency measures rsfMRI_E_ and DWI_E_ as well as nodal efficiency measures rsfMRI_Ei_ and DWI_Ei_. Sixth, global mediation analyses were performed for each combination of brain metric and PGS. Here, general intelligence as quantified by the I-S-T 2000 R sum score served as the dependent variable. Independent variables were one of the two PGS (PGS_EA_ and PGS_GI_). Whole brain measures (total surface area, mean cortical thickness, DWI_E_ or rsfMRI_E_) served as mediators. Finally, region-specific multi-mediator analyses were performed via elastic-net regression for each combination of brain metric and PGS. Again, the I-S-T 2000 R sum score was the dependent and PGS the independent variable. Surface area, cortical thickness, DWI_Ei_ or rsfMRI_Ei_ of each HCPMMP area served as mediators.

#### Analysis of diffusion-weighted data

Diffusion tensor modelling and probabilistic fiber tractography were conducted using the FDT toolbox (https://fsl.fmrib.ox.ac.uk/fsl/fslwiki/FDT) in FSL version 5.0.9. (https://fsl.fmrib.ox.ac.uk/fsl/fslwiki), following standard protocol (*54*). Image pre-processing included eddy currents correction and head motion correction. Additionally, the gradient directions of each volume were adjusted using the rotation parameters that were obtained from head motion correction. As described in previous section, the 180 cortical and 8 subcortical regions from each hemisphere were transformed into the native space of the diffusion-weighted images. Subsequently, these transformed regions were used as seed and target regions for probabilistic fiber tractography. To this end, we used a dual-fiber model implemented in the latest version of BEDPOSTX (https://users.fmrib.ox.ac.uk/~moisesf/Bedpostx_GPU/). This model allows for the representation of two fiber orientations per voxel and thus enables the modelling of crossing fibers, which produces more reliable results compared to single-fiber models (*55*). The classification targets approach implemented in FDT was used to perform probabilistic fiber tracking (*44*). Five thousand tract-following samples were generated at each voxel. The step length was 0.5 mm and the curvature threshold was 0.2 (only allowing for angles larger than 80 degrees). In order to quantify the connectivity between a seed voxel and a specific target region, the number of streamlines originating from the seed voxel and reaching the target region was determined. Subsequently, the overall connectivity between two brain regions was determined by calculating the sum of all streamlines proceeding from the seed to the target region and vice versa.

### Analysis of resting-state data

Resting-state data were pre-processed using the FSL toolbox MELODIC. The first two volumes of each resting-state scan were discarded. This was done to allow for signal equilibration, motion and slice timing correction, as well as high-pass temporal frequency filtering (0.005 Hz). We did not apply spatial smoothing to avoid the introduction of spurious correlations in neighboring voxels. Analogous to the analysis of the diffusion data, all brain regions were transformed into the native space of the resting-state images for functional connectivity analysis. For each region, a mean resting-state time course was calculated by averaging the time courses of all corresponding voxels. We computed partial correlations between the average time courses of all cortical and subcortical regions, while controlling for several nuisance variables, namely all six motion parameters as well as average time courses extracted from white matter regions and ventricles (*18*).

### Graph metrics

Graph metrics were calculated using the Brain Connectivity Toolbox (*56*) in combination with in-house MATLAB code. Each resting-state and DWI network consisted of 376 nodes, including 360 cortical (180 in each hemisphere) and 16 subcortical regions (8 in each hemisphere). We employed Holm-Bonferroni pruning with a threshold of 0 (a = 0.01, one tailed) as proposed by Ivković *et al*. (*57*) to remove spurious network connections. By following this approach, 65,357 edges from the DWI network and 748 edges from the resting-state network were removed. Two nodes (LH_H and RH_H) were removed from the resting-state network completely, as they did not show any connections to other nodes after pruning. Using the Brain Connectivity Toolbox, we computed global efficiency, a graph metric used in previous studies investigating the association between network connectivity and cognitive performance (*21, 28*). Global efficiency quantifies how efficiently information can be transferred across the brain (*58*). Large edge weights and small shortest path lengths typically lead to an increase in this metric. A shortest path is defined as the minimal number of edges it needs to connect a pair of nodes. The shortest path lengths between all pairs of nodes are comprised in the distance matrix *d*. This matrix can be created by calculating the inverse of the weighted adjacency matrix and running Dijksta’s algorithm (*59*). The global efficiency of one specific brain region is called nodal efficiency. It is calculated as the average inverse shortest path length between a node *i* and all other nodes *j* within a network *G* (*E_i_*). The global efficiency of the entire network is the average inverse shortest path length between each pair of nodes within *G (E)*:

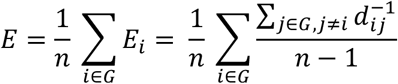

### Statistical analysis

All statistical analyses were conducted in R Studio (1.3.1093) with R version 4.1.0 (2021-05-18). Data points were treated as outliers if they deviated more than three interquartile ranges from the respective variable’s group mean (I-S-T 2000 R sum score, total surface area, mean cortical thickness, DWI or resting-state global efficiency). In such cases, all data from the corresponding participant were removed from analysis. No subjects were excluded from analyses concerning cortical surface area and cortical thickness (523 remaining subjects). Three subjects were excluded from analyses concerning DWI and resting-state network connectivity (520 remaining subjects).

#### Partial correlations

We computed partial correlations between the I-S-T 2000 R score, two PGS (EA and GI), and several brain parameters (total surface area, mean cortical thickness, DWI global efficiency and resting-state fMRI global efficiency) using the *partial.cor* function included in the *RcmdrMisc* package. Age and sex were treated as confounding variables and regressed out.

#### Global mediation model

We computed single mediation models using the *lavaan* package. In each model, the I-S-T 2000 R sum score served as the dependent variable, while one of the two PGS (EA and GI) served as the independent variable. For each model, we chose one out of four variables as the mediator, namely total surface area, mean cortical thickness, DWI global efficiency and resting-state fMRI global efficiency. Furthermore, we controlled for age, sex, and the first four principal components of the population stratification. We calculated one mediation for all PGS-mediator pairings, resulting in a total of eight mediations. Figure 1 (bottom half, middle box) shows a schematic depiction of a single mediation model. We used robust maximum likelihood estimator *MLM* with robust standard errors and a Satorra-Bentler scaled test statistic (*60*).

#### Brain area specific mediation via elastic-net regression

Following the computation of global mediation models, we investigated if a set of specific brain areas mediates the effect of PGS on I-S-T 2000 R. For this purpose, we employed *exploratory mediation analysis by regularization, a* tool developed to identify a subset of mediators from a large pool of potential mediators (*61, 62*). This approach does not use p-values to determine the statistical significance of a mediator. Hence, it does not require a standard correction procedure for multiple comparison (e.g. FDR or Bonferroni (*63*)). Instead, it utilizes regularization such as the least absolute shrinkage operator (lasso), which puts a penalty on effect sizes. Here, small effects are pushed down to zero and only strong effects remain non-zero. An in-depth explanation of this approach is provided by Serang *et al.* (*62*).

In short, all potential mediators are included in the model and the corresponding regression weights *a* and *b* are penalized (*64*). The penalty term lambda is determined using *k*-fold cross validation, which is a mechanism to prevent overfitting. Here, the data is split into *k* subsets. One of those subsets is selected as the testing-set while the rest of the data is used as the training-set. This is done *k* times with every subset being used as the testing-set once. The mediation effect of a mediator is calculated by multiplying the regression parameters *a* and *b*. If either parameter is regularized to zero, the mediation effect also becomes zero. If both *a* and *b* remain non-zero after regularization, the mediation effect will be non-zero as well. After this penalization procedure, all potential mediators with non-zero mediation effects are selected as mediators. While this method is a good way of eliminating mediators with small effect sizes, it also brings the effect sizes of real mediators close to zero. In order to address this potential bias, the model is fit again without penalization. With a model that only includes the pre-selected subset of mediators unbiased effect sizes can be acquired (*61*).

For the procedure described above, we used the *xmed* function from the *regsem* package (*61, 62, 64*). All variables were standardized and residualized for sex, age and the first four principal components of population stratification. The number of cross-validation folds was set to *k* = 80. The threshold for detecting non-zero mediation effects was set to 0.001 and the type of regression was set to elastic-net. Elastic-net is another type of regularized regression that combines lasso and ridge regression (*65*). The difference between ridge and lasso-regression is that the lasso penalty can shrink a parameter to zero, whereas ridge regression can only asymptotically shrink a parameter towards zero. Thus, lasso is suitable for models in which a lot of variables are expected to have no or little effect on the dependent variable, while ridge regression is suitable for models in which most variables are expected to have a considerable effect on the dependent variable. Elastic-net regression can be considered an ideal approach if one does not have clear expectations regarding every variable. In comparison to lasso regression, elastic-net regression is also better at handling correlations between variables (*65*), which was an important factor in our decision to choose elastic-net over lasso regression. Regularized elastic-net regression has already been successfully applied in a previous study investigating the association between fluid/crystallized intelligence and the microstructure of multiple white-matter tracts (*63*). We computed 8 mediator models for all combinations of PGS (GI, EA) and brain parameters (surface area, cortical thickness, DWI nodal-efficiency, resting-state fMRI nodal-efficiency). PGS served as the independent variable and the I-S-T 2000 R sum score as the dependent variable. After the pruning procedure described above (see Graph metrics), mediator models involving surface area and cortical thickness comprised 180 potential mediators each (180 cortical areas). The model involving DTI nodal-efficiency comprised 188 potential mediators (180 cortical and 8 subcortical areas) and the model involving resting-state nodal-efficiency comprised 187 potential mediators (179 cortical and 8 subcortical areas).

We employed an altered version of the function provided by Serang *et al.* (*61*). The modified code can be found at https://osf.io/2qamu/. First, we did not use the lambda sequence specified in the *xmed* function, but the default lambda sequence the *glmnet* package (*66*) which *xmed* interfaces to. This was done because the lambda sequence specified by *xmed* leads to ceiling effects in the lambda parameters. Second, we included the PGS as an independent variable in the regression of the mediators on the dependent variable. This effect was not penalized.

Apart from investigating which brain areas mediate the relationship between PGS and intelligence, we were also interested in the direct effects of PGS on the brain and the direct effects of the brain on intelligence (see Figure 1, paths *a* and *b*). Thus, we followed a similar approach to identify variables exhibiting non-zero effects within path *a* and path *b* regressions. The threshold for detecting non-zero effects was set to 0.01. This was done, because the mediation effects are the product of the regularized *a* and *b* parameters, which take values below 1. Hence, mediation effects are smaller compared to the regularized *a* and *b* parameters. Again, coefficients were re-estimated with *lavaan* to avoid biased effect sizes.

### Overlap of mediating areas and P-FIT

Finally, we aimed to test whether the mediating brain areas overlapped with the P-FIT network. It is important to note, that the P-FIT network is based on Brodmann areas (BA). In the original version proposed by Jung and Haier (*17*), the P-FIT features a network of 14 BA. In an updated version by Basten et al. (*67*) the network’s composition was confirmed, but also extended with 5 additional BA. In order to compare the HCPMMP areas from our analyses with P-FIT BA, we employed a cortical parcellation based on BA, which is included as an annotation file in FreeSurfer. This annotation file was converted to a volumetric segmentation matching the cortex of the fsaverage standard brain. The same was done to the HCPMMP annotation file. By means of an in-house MATLAB program, the overlap between all HCPMMP and BA areas was calculated. An HCPMMP area was specified as being part of the P-FIT network when it showed at least 80% overlap with one or more P-FIT BA in both hemispheres. This was true for 87 HCPMMP areas (Table S4).

## Results

In preliminary analyses, to gain an overview of bivariate correlations and to compare our data with previously reported results, partial correlations were computed to test the associations between PGS and intelligence, PGS and whole brain properties, as well as whole brain properties and intelligence (see Table 1). Both PGS were significantly associated with the I-S-T 2000 R sum score (see Table 1) and total surface area. PGS_EA_ was also associated with DWI_E_. The I-S-T 2000 R sum score was associated with both total surface area and DWI_E_. Mean cortical thickness and rsfMRI_E_ were not associated with PGS or the I-S-T 2000 sum score (see Table 1).

**Table 1.**
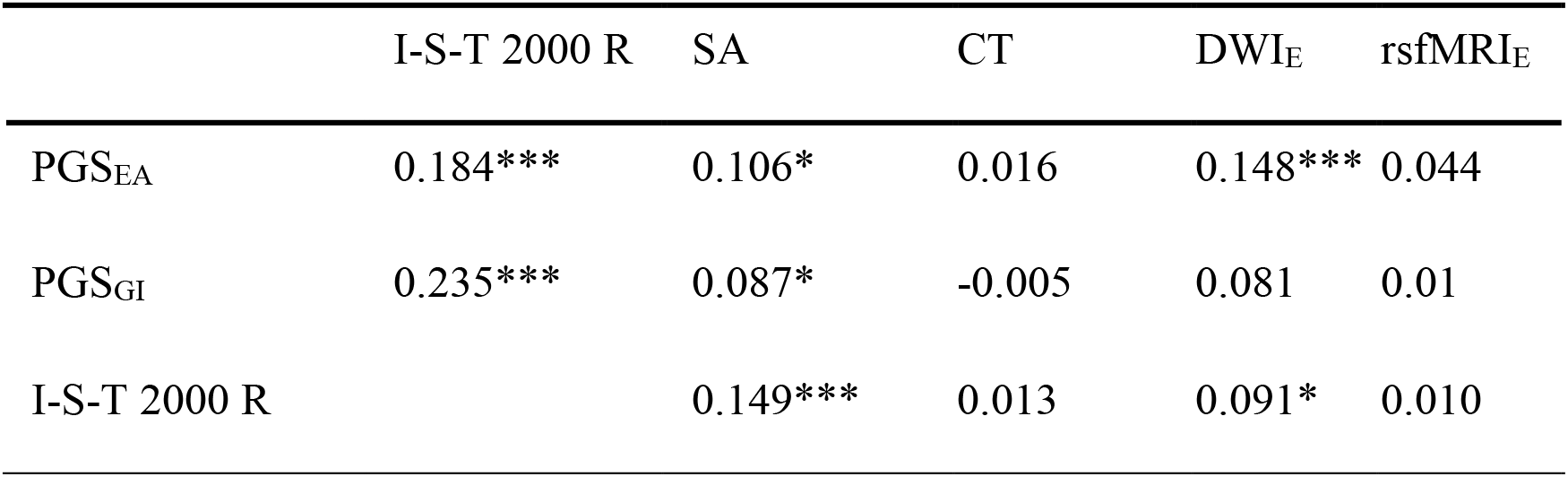
Partial correlation. Partial correlation coefficients (Pearson’s *r*) between I-S-T 2000 R performance, PGS for education attainment (EA), general intelligence (GI) and brain properties. Age and sex were used as controlling variables. SA = surface Area, CT = cortical thickness, DWI_E_ = DWI-network global efficiency, rsfMRI_E_ = resting-state network global efficiency. * *p* < 0.05, ** *p* < 0.01, *** *p* < 0.001 (two-tailed).

### Global mediation analysis

Results of the global mediation analysis are shown in Figure 2. PGS_EA_ was significantly associated with total surface area and DWI_E_. Total surface area and DWI_E_ were significantly associated with the I-S-T 2000 sum score. However, none of the brain parameters turned out to be significant mediators in the effect of PGS on general intelligence on a whole brain level (all *p* > .08).

**Figure 2.**
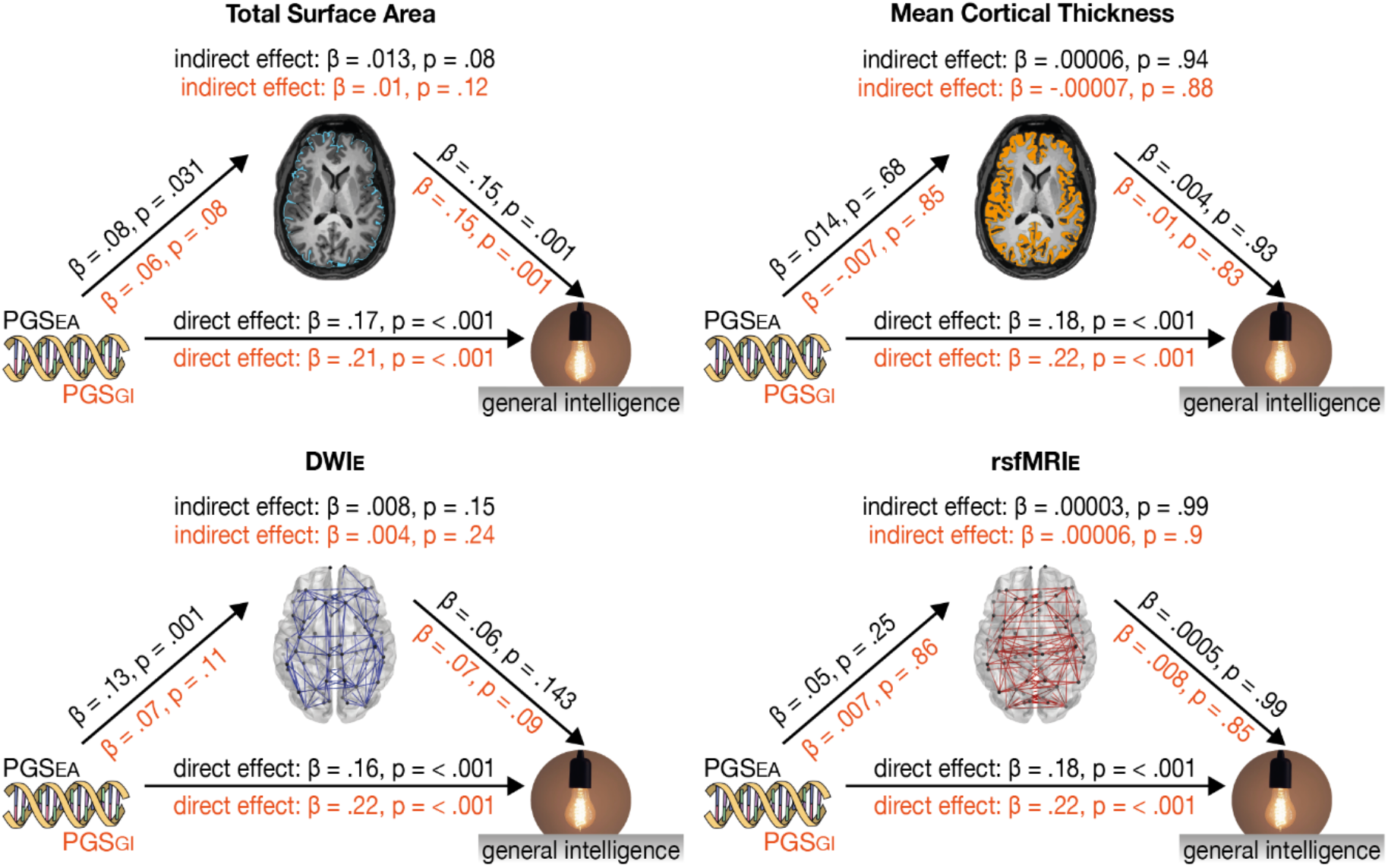
Results of the global mediation analysis. We used total surface area, mean cortical thickness, DWI_E_ and rsfMRI_E_ as mediators. In all cases, general intelligence, as measured by the I-S-T 2000 R sum score, served as the dependent variable. PGS_EA_ or PGS_GI_ served as independent variables. Effect sizes and p-values are depicted in black (above the arrows) for analyses with PGS_EA_ and in orange (below the arrows) for analyses with PGS_GI_.

### Brain area specific mediation

#### Surface area

Results of the region-specific multi-mediator analysis via elastic net showed that PGS_EA_ was associated with the surface area of the majority of HCPMMP areas (112 areas; Figure 3). All effects except one were positive, indicating that higher PGS_EA_ is associated with larger surface area (path *a*). Furthermore, the surface area of 18 brain areas in the parietal and frontal cortices was associated with the I-S-T 2000 R sum score (path *b*). Ten out of these brain areas mediated the effects of PGS_EA_ on general intelligence (*a*b*). HCPMMP areas 4 (primary motor cortex), 6r (premotor cortex), MIP, IP1 (intraparietal areas), OFC (orbital frontal cortex), OP1 (parietal operculum), STGa (anterior superior temporal gyrus), and PH (posterior temporal cortex) showed positive mediation effects. HCPMMP area 1 (somatosensory cortex) and IFSa (inferior frontal sulcus) showed negative mediation effects. Half of these areas was found to be part of the P-FIT network (MIP, 6r, IFSa, PH, IP1).

**Figure 3.**
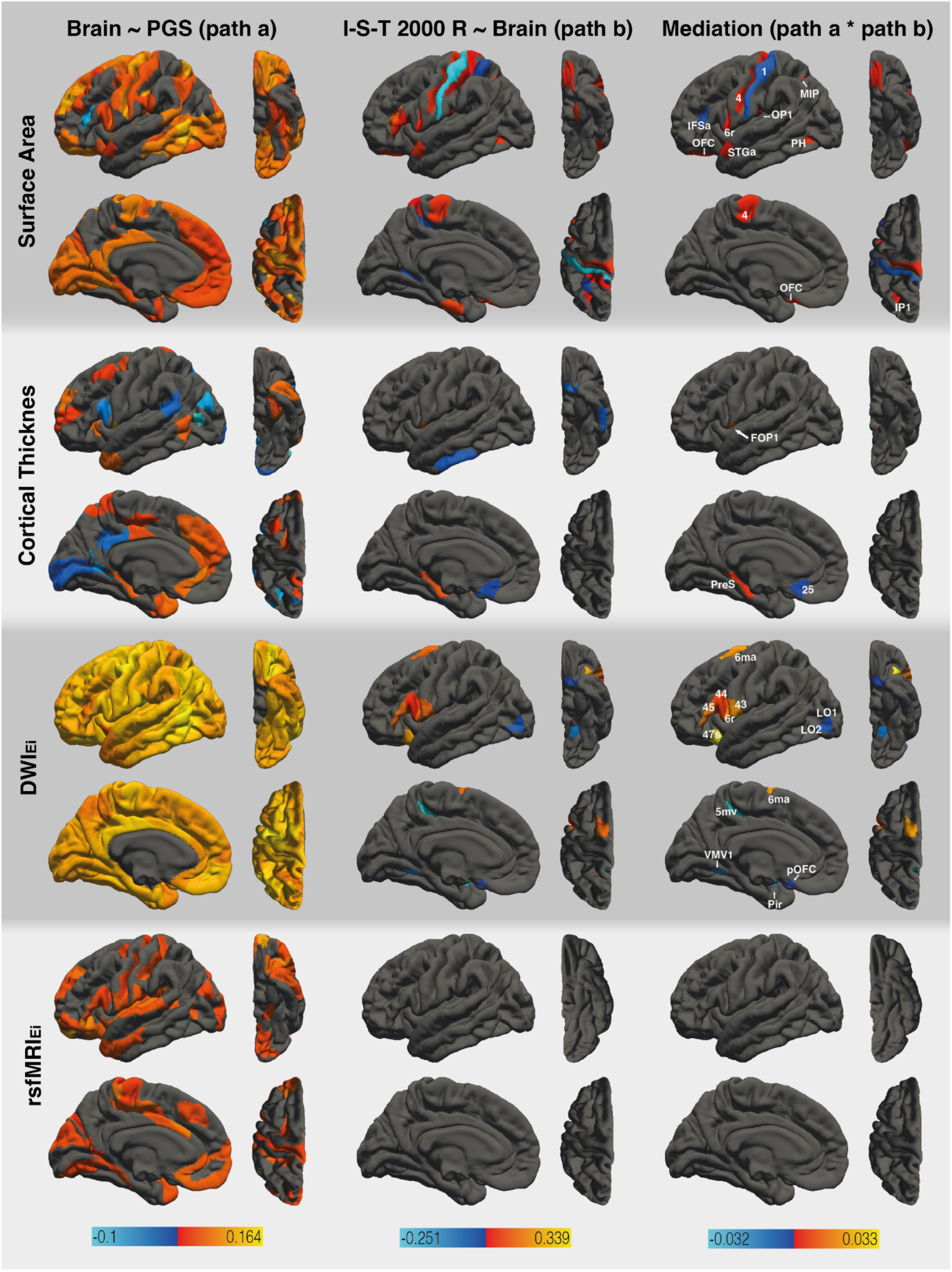
Results of the region-specific multi-mediator analysis via elastic net. with PGS_EA_ as dependent variable. The analysis employed the following mediators: surface area, cortical thickness, DWI_Ei_, and rsfMRI_Ei_ (from top to bottom). The figure shows the results from path *a* analysis, path *b* analysis, and the mediation effect (from left to right). Brain surfaces are shown in lateral, inferior, sagittal, and superior view (from left to right). Positive effects are depicted in red and yellow, negative effects are depicted in blue. Colored mediating areas are labeled according to the HCPMMP. Path *a* analysis of DWI_Ei_ also revealed positive associations between PGS_EA_ and eight subcortical areas. For a full list of areas and effect sizes see Tables S2 and S3.

Similar results were obtained when PGS_GI_ was used as the predictor (see Figure 4). We found PGS_GI_ to be associated with the surface area of 87 brain areas distributed all over the cortex, with most areas largely matching (83%) those identified by the PGS_EA_ analysis. The surface area of eight areas was associated with general intelligence. Three of these areas mediated the effects of PGS_GI_ on general intelligence, namely HCPMMP areas MIP, IP1 (intraparietal areas), and PH (posterior temporal cortex). It is noteworthy, that all of these areas were identified as mediators in the PGS_EA_ analysis as well. All areas were found to be part of the P-FIT network.

**Figure 4.**
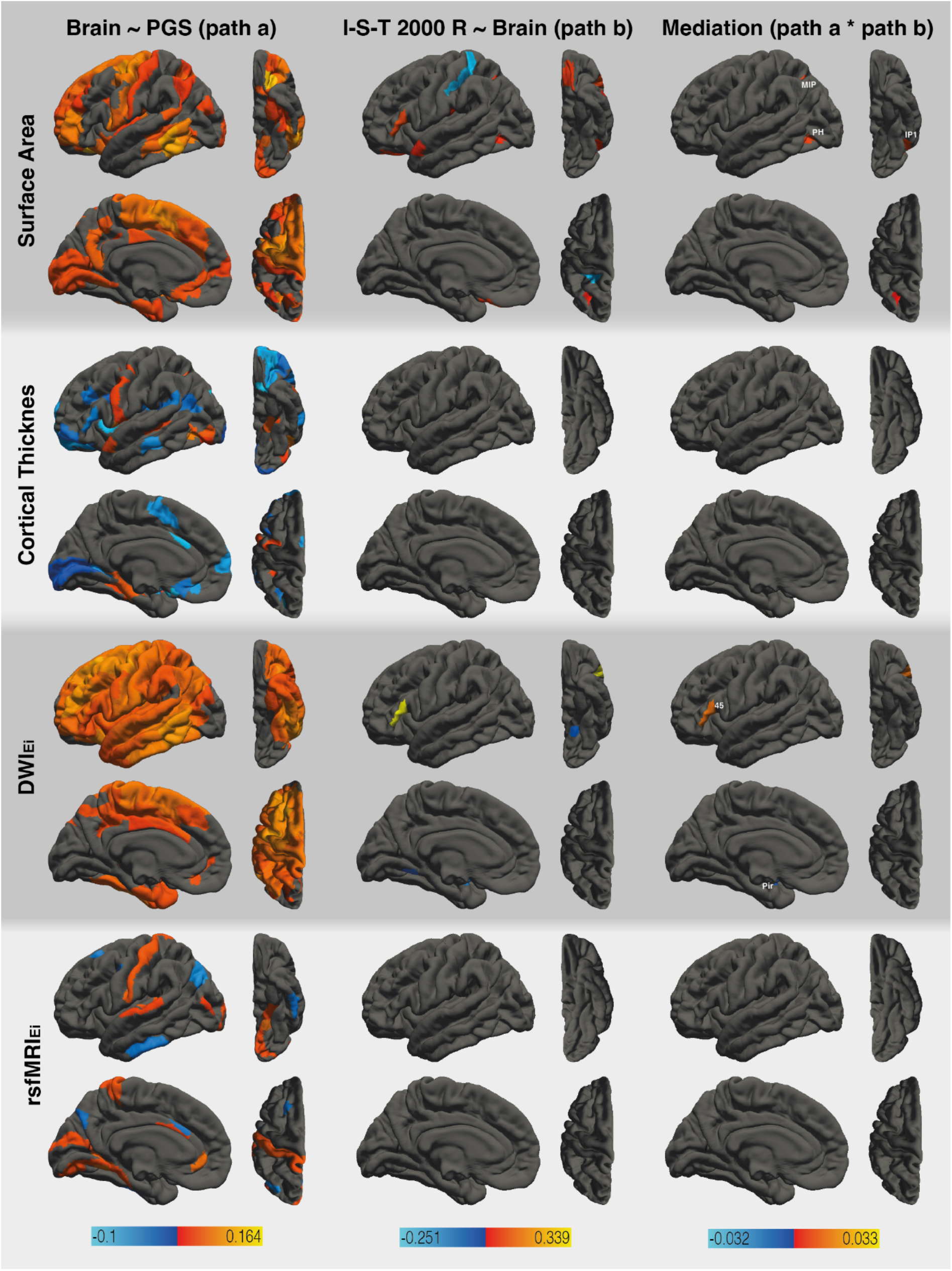
Results of the region-specific multi-mediator analysis via elastic net with PGS_GI_ as dependent variable. The analysis employed the following mediators: surface area, cortical thickness, DWI_Ei_, and rsfMRI_Ei_ (from top to bottom). The figure shows the results from path *a* analysis, path *b* analysis, and the mediation effect (from left to right). Brain surfaces are shown in lateral, inferior, sagittal, and superior view (from left to right). Positive effects are depicted in red and yellow, negative effects are depicted in blue. Colored mediating areas are labeled according to the HCPMMP. Path *a* analysis of DWI_Ei_ also revealed positive associations between PGS_GI_ and six subcortical areas. For a full list of areas and effect sizes see Tables S2 and S3.

#### Cortical thickness

PGS_EA_ was associated with cortical thickness in 39 brain areas, of which 26 (67%) exhibited positive effects and 13 (33%) exhibited negative effects. Seven cortical areas showed significant associations between cortical thickness and general intelligence. Three of these areas mediated the effects of PGS_EA_ on the I-S-T 2000 R sum score. HCPMMP areas FOP1 (frontal operculare area) and PreS (presubiculum) exhibited positive mediation effects, while area 25 (superior anterior cingulate cortex) exhibited a negative mediation effect. PGS_GI_ was associated with cortical thickness in 48 areas (21 positive associations and 27 negative associations) with limited overlap (23%) between the PGS_EA_ and PGS_GI_ analyses. The PGS_GI_ mediation model did not yield any areas which showed significant associations between cortical thickness and general intelligence. Consequently, no mediators for the effects of PGS_GI_ on general intelligence could be identified. None of the mediating areas were part of the P-FIT network.

#### DWI network efficiency

PGS_EA_ was positively associated with DWI_Ei_ in all cortical and subcortical areas (188). There were 12 areas in which DWI_E_ was associated with general intelligence. All of these areas were also mediators regarding the effects of PGS_EA_ on general intelligence. HCPMMP areas 6ma (anterior supplementary motor cortex), 6r (premotor cortex), 44, 45 (inferior frontal gyrus), 47s (orbitofrontal cortex), and 43 (posterior opercular cortex) exhibited positive mediation effects. HCPMMP areas LO1, LO2 (lateral occipital cortex), 5mv (superior parietal cortex), VMV1 (ventromedial visual area), Pir (piriform cortex), and pOFC (posterior orbitofrontal complex) exhibited negative mediation effects. Half of these areas was found to be part of the P-FIT network (LO1, LO2, 44, 45, 6r, 6ma). Similarly, PGS_GI_ was positively associated with DWI_E_ in 144 areas. DWI_E_ was associated with general intelligence in three areas and two of them, namely HCPMMP areas 45 (inferior frontal gyrus) and Pir (piriform cortex), were also found to be mediators regarding the effects of PGS_GI_ on general intelligence. These two areas were also identified as mediators in the PGS_EA_ analysis. HCPMMP area 45 was found to be part of the P-FIT network.

#### rsfMRI network efficiency

PGS_EA_ was associated with rsfMRI_E_ in 68 areas and all of these associations were positive. There were no areas which exhibited significant associations between rsfMRI_E_ and general intelligence or mediated the effects of PGS_EA_ on general intelligence. PGS_GI_ was associated with rsfMRI_E_ in 25 areas, with 19 (76%) of them showing positive associations. There were no areas which exhibited significant associations between rsfMRI_E_ and general intelligence or mediated the effects of PGS_GI_ on general intelligence.

Complete lists of HCPMMP areas and effect sizes can be found in Tables S2 (path *a*), S3 (path *b*), and S4 (mediation).

## Discussion

Genetic variability robustly predicts interindividual differences in intelligence, but it is still largely unknown which neurobiological intermediates are involved in the path form genetic disposition to phenotype. Hence, it was the aim of our study to conduct integrative analyses encompassing genome-wide SNP variability, in-depth brain imaging, and a detailed measurement of cognitive abilities. By doing so, we were able to show that regional surface area and structural network efficiency are mediators of the relationship between genetic disposition and measured intelligence.

In line with other studies, PGS significantly predicted cognitive abilities. Furthermore, PGS were associated with morphological and connectivity brain measures of widely distributed cortical and subcortical regions, a finding which is in accordance with previously reported results showing genetic correlations between cognitive abilities and brain structure (*33*). To further investigate which of these brain areas link genetic variation to differences in cognitive abilities, four brain properties on global and regional level were tested as putative mediators. rsfMRI was not associated with cognitive abilities, neither on a global scale nor on the level of brain regions. In case of cortical thickness, there was limited evidence of mediation effects. However, the surface area and structural connectivity of several brain areas were associated with intelligence and also identified as mediators.

With regard to surface area, we found ten brain regions that mediated the effects of PGS_EA_ on general intelligence. Respective areas were mainly located in the posterior parietal, posterior temporal, and superior frontal cortices. Three of these areas were also identified when PGS_GI_ was used as predictor and half of the mediating areas were part of the P-FIT network (MIP, 6r, IFSa, PH, IP1). There were five brain areas outside of the P-FIT network, namely the primary motor cortex (4), the primary somatosensory cortex (1), the orbitofrontal cortex (OFC), and the posterior part of the parietal operculum (OP1). The common observation that the volume or surface area of cortical gray matter is positively associated with intelligence is typically explained in the following way. Individuals with more cortical volume or surface area are likely to possess more neurons (*68, 69*). A higher count in cortical neurons also indicates a higher number of synapses (*70*). Therefore, it is assumed that individuals with more cortical gray matter have more computational power to engage in problem solving and logical reasoning (*45*). Following this explanation, our results indicate that the SNPs associated with cognitive abilities may influence the gene expression related to neuron and synapse count within specific cortical areas. This in turn might influence intelligent thinking.

Our findings related to non-P-FIT areas are largely in line with the findings by Lett *et al.* (*42*), who also found a mediating effect of surface area in parts of the primary motor cortex, the orbitofrontal cortex, and the parietal operculum. The orbitofrontal cortex and its interaction with the anterior cingulate cortex have been associated with decision making (*71*). The orbitofrontal cortex encodes the value of available choices based on past experiences. The anterior cingulate cortex is involved in a more “down-stream” processing of decision consequences (*72*). While the primary motor cortex is usually not associated with intelligence, its structural and functional properties have been found to change in accordance with verbal and non-verbal intelligence in teenagers (*73*). The authors argue that this finding is indicative of an interrelation between cognitive and motor development (*72*), which may also be one reason behind the association between motor skills, cognitive performance, and academic achievements (*74, 75*). Although the primary somatosensory cortex was not identified as a mediator by Lett *et al.* (*42*) or Elliot *et al.* (*41*), a meta-analysis revealed its functional properties to be associated with fluid intelligence (*76*).

Many biological theories of intelligence highlight the importance of efficient information exchange across the brain. Naturally, this task is heavily dependent on the structural quality of an extensive brain network. A neuronal circuitry associated with higher intelligence is thought to foster a more directed information processing along relevant areas within the network. Our findings support this assumption by showing that the structural nodal efficiency of twelve brain areas mediated the relationship between PGS_EA_ and general intelligence. Moreover, two of these brain areas were also identified in the PGS_GI_ analyses. It is noteworthy that half of the mediating areas are part of the P-FIT network (6ma, 6r, 44, 45, LO1, LO2). The inferior frontal gyrus (45, 44), the premotor cortex (6r), and the anterior supplementary motor cortex (6ma) exhibited positive mediation effects. Parts of the lateral orbitofrontal cortex (LO1, LO2) exhibited negative mediating effects, which was due to negative associations between their structural connectivity and general intelligence. We also observed multiple mediators outside of the P-FIT network. The ventromedial visual area (VMV1) and posterior orbitofrontal cortex (pOFC) exhibited negative mediation effects, which was due to negative associations between their nodal efficiency and general intelligence. The orbitofrontal cortex (47s) and posterior opercular cortex (43) exhibited positive mediation effects. Functional properties of the right orbitofrontal cortex have been shown to be positively associated with fluid intelligence in a recent meta-analysis (*76*). The posterior opercular cortex is part of the so called cingulo-opercular network (*77*) which plays a critical role for intelligence according to the Network Neuroscience Theory (*12*). This theory proposes that the neural basis of general intelligence is manifested in the dynamics of multiple brain wide modular networks. In other words, the Network Neuroscience Theory emphasizes that intelligence depends on the efficiency with which specific brain networks can be reorganized and adapted to a situation. It has to be noted that this theory is focused on the dynamic state of networks and largely based on functional studies. Hence, it may not directly be applicable to white matter connectivity, even though functional networks have been proposed to arise from structural connectivity (*78*). The Network Neuroscience Theory proposes that crystallized intelligence relies on easy-to-reach functional network states which in turn rely on strong connections between some highly connected brain areas. In contrast, fluid intelligence is supposed to rely on difficult-to-reach network states, which in turn rely on weak connections between networks. Weak connections giving rise to difficult-to-reach network states are located in the frontoparietal network and the cingulo-opercular network (*12*). For the most part, P-FIT emerged from macrostructural studies. When looking at intelligence from a connectivity-based perspective, as is done in Network Neuroscience Theory, it seems plausible that there are brain areas whose morphological properties are not related to intelligence, while their connectivity patterns are. Our results support this assumption by showing that a group of SNPs, identified by GWAS, is likely to influence the gene expression shaping the structural efficiency of specific areas from an extensive and intelligence-related brain network.

Our results concerning surface area and structural connectivity show that there are considerably more brain areas mediating the effect between PGS_EA_ and general intelligence than between PGS_GI_ and general intelligence. In all likelihood, this is due to the difference in discovery sample sizes of respective GWAS. PGS_EA_ was derived from a GWAS with a sample size of 1,131,881 individuals (*10*), whereas PGS_GI_ was derived from a GWAS with a sample size of 269,867 individuals (*6*). This results in greater predictive power of PGS_EA_. While PGS_GI_ exhibited a stronger association with general intelligence in our sample compared to PGS_EA_, PGS_EA_ exhibited stronger associations with the analyzed brain properties (see Table 1). Some of the genes identified by Lee *et al.* (*10*) (PGS_EA_) are highly expressed in the brain prenatally and thus influence very early stages of brain development. Other genes show high expression both prenatally and postnatally. Functionally, the identified genes are involved in neurotransmitter secretion, the activation of ion channels and metabotropic receptors, as well as synaptic plasticity. Importantly, these genes are expressed in all parts of the nervous system and not limited to a certain set of brain areas. Our results are in line with this finding given that PGS_EA_ were associated with brain properties all over the cortex (Figure 3). However, since our analyses also included the phenotype, we were able to specify which parts of the brain are affected by intelligence-related gene expression as identified by Lee *et al.* (*10*). Importantly, this approach goes one step beyond investigating the genetic correlation between cognitive and brain phenotypes.

Whereas the direction of effect from genes to cognitive abilities and genes to brain structure is causal by definition (*5*), it is conceivable that there is a bidirectional relationship between brain structure and cognitive abilities. Recent analyses used bidirectional latent causal variable and Mendelian randomization to assess the causal direction between human cortical structure, general intelligence, and educational attainment. They provide evidence for an influence of brain structure on general intelligence and educational attainment (*33*). Our investigation includes both measured brain phenotype and detailed characterization of cognitive abilities and thus provides further evidence of causal processes between genetic variability and cognition through variation in brain structure and network connectivity.

There are certain limitations to our study. First, PGS for educational attainment tend to overestimate genetically caused effects in non-related samples (*10, 79*). Lee *et al.* (*10*) showed that the predictive power of PGS declines as much as 40% when within-family differences in educational attainment are taken into account, which is partly due to gene-environment correlations. The genes of parents also influence the rearing environment of their child, which results in a correlation between the environment and the genes a child inherits from their parents (*79*). The effect of parental genes on rearing environment is demonstrated by the observation that even non-shared genetic information of parents are predictive of a child’s educational attainment (*80*). Thus, the predictive power of the PGS utilized in our study can in part be attributed to gene-environment correlations. Second, our functional connectivity analysis did not identify any regions that mediated the effects of PGS on general intelligence; or any brain area that was directly associated with general intelligence (see Figures 3 and 4). In order to compute nodal efficiency, we aggregated resting-state data across the entire time span of our recordings. However, Network Neuroscience Theory argues that the crucial aspect of intelligence-related functional networks is their dynamic flexibility (*12*), which is not captured by the metrics we used. Hence, it is indeed conceivable that the flexibility of specific networks mediates the effects of genetic variation on general intelligence. Future studies using temporally high resolution rsfMRI and dynamic connectivity analyses should investigate the mediation effects of dynamic connectivity metrics.

This study is the first to investigate the mediating effects of multimodal, region-specific brain properties on the association between genetic variation and intelligence. We show that the surface area and structural connectivity of frontal, sensory, motor, temporal, and anterior occipital brain regions provide a missing piece in the link between genetic variation and general intelligence. These findings are a crucial step forward in decoding the neurogenetic underpinnings of intelligence, as they identify specific regional networks that relate polygenic variation to intelligence.

## Acknowledgements

The authors would like to thank all research assistants for their support during the behavioral measurements.

## Data and materials availability

The data and R code that support the findings of this study are available from the corresponding author upon reasonable request or can be downloaded from an Open Science Framework repository [https://osf.io/2qamu/].

## References

1. I. J. Deary, S. R. Cox, W. D. Hill, Genetic variation, brain, and intelligence differences. Mol Psychiatry (2021).

2. I. J. Deary, The Stability of Intelligence From Childhood to Old Age. Curr Dir Psychol Sci 23, 239–245 (2014).

3. I. J. Deary, S. Strand, P. Smith, C. Fernandes, Intelligence and educational achievement. Intelligence 35, 13–21 (2007).

4. C. M. Calvin, G. D. Batty, G. Der, C. E. Brett, A. Taylor, A. Pattie, I. Čukić, I. J. Deary, Childhood intelligence in relation to major causes of death in 68 year follow-up: prospective population study. BMJ 357, 1–14 (2017).

5. R. Plomin, S. von Stumm, The new genetics of intelligence. Nat Rev Genet 19, 148–159 (2018).

6. J. E. Savage, P. R. Jansen, S. Stringer, K. Watanabe, J. Bryois, C. A. de Leeuw, M. Nagel, S. Awasthi, P. B. Barr, J. R. I. Coleman, K. L. Grasby, A. R. Hammerschlag, J. A. Kaminski, R. Karlsson, E. Krapohl, M. Lam, M. Nygaard, C. A. Reynolds, J. W. Trampush, H. Young, D. Zabaneh, S. Hägg, N. K. Hansell, I. K. Karlsson, S. Linnarsson, G. W. Montgomery, A. B. Muñoz-Manchado, E. B. Quinlan, G. Schumann, N. G. Skene, B. T. Webb, T. White, D. E. Arking, D. Avramopoulos, R. M. Bilder, P. Bitsios, K. E. Burdick, T. D. Cannon, O. Chiba-Falek, A. Christoforou, E. T. Cirulli, E. Congdon, A. Corvin, G. Davies, I. J. Deary, P. DeRosse, D. Dickinson, S. Djurovic, G. Donohoe, E. D. Conley, J. G. Eriksson, T. Espeseth, N. A. Freimer, S. Giakoumaki, I. Giegling, M. Gill, D. C. Glahn, A. R. Hariri, A. Hatzimanolis, M. C. Keller, E. Knowles, D. Koltai, B. Konte, J. Lahti, S. Le Hellard, T. Lencz, D. C. Liewald, E. London, A. J. Lundervold, A. K. Malhotra, I. Melle, D. Morris, A. C. Need, W. Ollier, A. Palotie, A. Payton, N. Pendleton, R. A. Poldrack, K. Räikkönen, I. Reinvang, P. Roussos, D. Rujescu, F. W. Sabb, M. A. Scult, O. B. Smeland, N. Smyrnis, J. M. Starr, V. M. Steen, N. C. Stefanis, R. E. Straub, K. Sundet, H. Tiemeier, A. N. Voineskos, D. R. Weinberger, E. Widen, J. Yu, G. Abecasis, O. A. Andreassen, G. Breen, L. Christiansen, B. Debrabant, D. M. Dick, A. Heinz, J. Hjerling-Leffler, M. A. Ikram, K. S. Kendler, N. G. Martin, S. E. Medland, N. L. Pedersen, R. Plomin, T. J. C. Polderman, S. Ripke, S. van der Sluis, P. F. Sullivan, S. I. Vrieze, M. J. Wright, D. Posthuma, Genome-wide association meta-analysis in 269,867 individuals identifies new genetic and functional links to intelligence. Nat Genet 50, 912–919 (2018).

7. S. W. Choi, T. S.-H. Mak, P. F. O’Reilly, Tutorial: a guide to performing polygenic risk score analyses. Nat Protoc 15, 2759–2772 (2020).

8. F. Dudbridge, Power and predictive accuracy of polygenic risk scores. PLoS Genet 9, e1003348 (2013).

9. D. Dima, G. Breen, Polygenic risk scores in imaging genetics: Usefulness and applications. J Psychopharmacol 29, 867–871 (2015).

10. J. J. Lee, R. Wedow, A. Okbay, E. Kong, O. Maghzian, M. Zacher, T. A. Nguyen-Viet, P. Bowers, J. Sidorenko, R. Karlsson Linnér, M. A. Fontana, T. Kundu, C. Lee, H. Li, R. Li, R. Royer, P. N. Timshel, R. K. Walters, E. A. Willoughby, L. Yengo, M. Alver, Y. Bao, D. W. Clark, F. R. Day, N. A. Furlotte, P. K. Joshi, K. E. Kemper, A. Kleinman, C. Langenberg, R. Mägi, J. W. Trampush, S. S. Verma, Y. Wu, M. Lam, J. H. Zhao, Z. Zheng, J. D. Boardman, H. Campbell, J. Freese, K. M. Harris, C. Hayward, P. Herd, M. Kumari, T. Lencz, J. Luan, A. K. Malhotra, A. Metspalu, L. Milani, K. K. Ong, J. R. B. Perry, D. J. Porteous, M. D. Ritchie, M. C. Smart, B. H. Smith, J. Y. Tung, N. J. Wareham, J. F. Wilson, J. P. Beauchamp, D. C. Conley, T. Esko, S. F. Lehrer, P. K. E. Magnusson, S. Oskarsson, T. H. Pers, M. R. Robinson, K. Thom, C. Watson, C. F. Chabris, M. N. Meyer, D. I. Laibson, J. Yang, M. Johannesson, P. D. Koellinger, P. Turley, P. M. Visscher, D. J. Benjamin, D. Cesarini, Gene discovery and polygenic prediction from a genome-wide association study of educational attainment in 1.1 million individuals. Nat Genet 50, 1112–1121 (2018).

11. I. J. Deary, L. Penke, W. Johnson, The neuroscience of human intelligence differences. Nat Rev Neurosci 11, 201–211 (2010).

12. A. K. Barbey, Network Neuroscience Theory of Human Intelligence. Trends Cogn Sci 22, 8–20 (2018).

13. J. Pietschnig, L. Penke, J. M. Wicherts, M. Zeiler, M. Voracek, Meta-analysis of associations between human brain volume and intelligence differences: How strong are they and what do they mean? Neurosci Biobehav Rev 57, 411–432 (2015).

14. M. McDaniel, Big-brained people are smarter: A meta-analysis of the relationship between in vivo brain volume and intelligence. Intelligence 33, 337–346 (2005).

15. K. L. Narr, R. P. Woods, P. M. Thompson, P. Szeszko, D. Robinson, T. Dimtcheva, M. Gurbani, A. W. Toga, R. M. Bilder, Relationships between IQ and regional cortical gray matter thickness in healthy adults. Cereb Cortex 17, 2163–2171 (2007).

16. Y. Y. Choi, N. A. Shamosh, S. H. Cho, C. G. DeYoung, M. J. Lee, J.-M. Lee, S. I. Kim, Z.-H. Cho, K. Kim, J. R. Gray, K. H. Lee, Multiple bases of human intelligence revealed by cortical thickness and neural activation. J Neurosci 28, 10323–10329 (2008).

17. R. E. Jung, R. J. Haier, The Parieto-Frontal Integration Theory (P-FIT) of intelligence: converging neuroimaging evidence. Behav Brain Sci 30, 135–54; discussion 154-87 (2007).

18. C. Fraenz, C. Schlüter, P. Friedrich, R. E. Jung, O. Güntürkün, E. Genç, Interindividual differences in matrix reasoning are linked to functional connectivity between brain regions nominated by Parieto-Frontal Integration Theory. Intelligence 87, 101545 (2021).

19. Y. Li, Y. Liu, J. Li, W. Qin, K. Li, C. Yu, T. Jiang, Brain anatomical network and intelligence. PLoS Comput Biol 5, e1000395 (2009).

20. J. A. Pineda-Pardo, K. Martínez, F. J. Román, R. Colom, Structural efficiency within a parieto-frontal network and cognitive differences. Intelligence 54, 105–116 (2016).

21. J. Ma, H. J. Kang, J. Y. Kim, H. S. Jeong, J. J. Im, E. Namgung, M. J. Kim, S. Lee, T. D. Kim, J. K. Oh, Y.-A. Chung, I. K. Lyoo, S. M. Lim, S. Yoon, Network attributes underlying intellectual giftedness in the developing brain. Sci Rep 7, 11321 (2017).

22. F. U. Fischer, D. Wolf, A. Scheurich, A. Fellgiebel, Association of structural global brain network properties with intelligence in normal aging. PloS one 9, e86258 (2014).

23. D.-J. Kim, E. P. Davis, C. A. Sandman, O. Sporns, B. F. O’Donnell, C. Buss, W. P. Hetrick, Children’s intellectual ability is associated with structural network integrity. NeuroImage 124, 550–556 (2016).

24. W. Wen, W. Zhu, Y. He, N. A. Kochan, S. Reppermund, M. J. Slavin, H. Brodaty, J. Crawford, A. Xia, P. Sachdev, Discrete neuroanatomical networks are associated with specific cognitive abilities in old age. J. Neurosci. 31, 1204–1212 (2011).

25. S. J. Wiseman, T. Booth, S. J. Ritchie, S. R. Cox, S. Muñoz Maniega, M. C. Del Valdés Hernández, D. A. Dickie, N. A. Royle, J. M. Starr, I. J. Deary, J. M. Wardlaw, M. E. Bastin, Cognitive abilities, brain white matter hyperintensity volume, and structural network connectivity in older age. Hum Brain Mapp 39, 622–632 (2018).

26. M. D. Fox, M. E. Raichle, Spontaneous fluctuations in brain activity observed with functional magnetic resonance imaging. Nat Rev Neurosci 8, 700–711 (2007).

27. M. P. van den Heuvel, C. J. Stam, R. S. Kahn, H. E. Hulshoff Pol, Efficiency of functional brain networks and intellectual performance. J Neurosci 29, 7619–7624 (2009).

28. J. D. Kruschwitz, L. Waller, L. S. Daedelow, H. Walter, I. M. Veer, General, crystallized and fluid intelligence are not associated with functional global network efficiency: A replication study with the human connectome project 1200 data set. NeuroImage 171, 323–331 (2018).

29. K. Hilger, M. Ekman, C. J. Fiebach, U. Basten, Intelligence is associated with the modular structure of intrinsic brain networks. Sci Rep 7, 16088 (2017).

30. K. Hilger, M. Ekman, C. J. Fiebach, U. Basten, Efficient hubs in the intelligent brain: Nodal efficiency of hub regions in the salience network is associated with general intelligence. Intelligence 60, 10–25 (2017).

31. B. Zhao, T. Li, Y. Yang, X. Wang, T. Luo, Y. Shan, Z. Zhu, Di Xiong, M. E. Hauberg, J. Bendl, J. F. Fullard, P. Roussos, Y. Li, J. L. Stein, H. Zhu, Common genetic variation influencing human white matter microstructure. Science 372, eabf3736 (2021).

32. B. Zhao, J. Zhang, J. G. Ibrahim, T. Luo, R. C. Santelli, Y. Li, T. Li, Y. Shan, Z. Zhu, F. Zhou, H. Liao, T. E. Nichols, H. Zhu, Large-scale GWAS reveals genetic architecture of brain white matter microstructure and genetic overlap with cognitive and mental health traits (n = 17,706). Mol Psychiatry 26, 3943–3955 (2021).

33. K. L. Grasby, N. Jahanshad, J. N. Painter, L. Colodro-Conde, J. Bralten, D. P. Hibar, P. A. Lind, F. Pizzagalli, C. R. K. Ching, M. A. B. McMahon, N. Shatokhina, L. C. P. Zsembik, S. I. Thomopoulos, A. H. Zhu, L. T. Strike, I. Agartz, S. Alhusaini, M. A. A. Almeida, D. Alnæs, I. K. Amlien, M. Andersson, T. Ard, N. J. Armstrong, A. Ashley-Koch, J. R. Atkins, M. Bernard, R. M. Brouwer, E. E. L. Buimer, R. Bülow, C. Bürger, D. M. Cannon, M. Chakravarty, Q. Chen, J. W. Cheung, B. Couvy-Duchesne, A. M. Dale, S. Dalvie, T. K. de Araujo, G. I. de Zubicaray, S. M. C. de Zwarte, A. den Braber, N. T. Doan, K. Dohm, S. Ehrlich, H.-R. Engelbrecht, S. Erk, C. C. Fan, I. O. Fedko, S. F. Foley, J. M. Ford, M. Fukunaga, M. E. Garrett, T. Ge, S. Giddaluru, A. L. Goldman, M. J. Green, N. A. Groenewold, D. Grotegerd, T. P. Gurholt, B. A. Gutman, N. K. Hansell, M. A. Harris, M. B. Harrison, C. C. Haswell, M. Hauser, S. Herms, D. J. Heslenfeld, N. F. Ho, D. Hoehn, P. Hoffmann, L. Holleran, M. Hoogman, J.-J. Hottenga, M. Ikeda, D. Janowitz, I. E. Jansen, T. Jia, C. Jockwitz, R. Kanai, S. Karama, D. Kasperaviciute, T. Kaufmann, S. Kelly, M. Kikuchi, M. Klein, M. Knapp, A. R. Knodt, B. Krämer, M. Lam, T. M. Lancaster, P. H. Lee, T. A. Lett, L. B. Lewis, I. Lopes-Cendes, M. Luciano, F. Macciardi, A. F. Marquand, S. R. Mathias, T. R. Melzer, Y. Milaneschi, N. Mirza-Schreiber, J. C. V. Moreira, T. W. Mühleisen, B. Müller-Myhsok, P. Najt, S. Nakahara, K. Nho, L. M. Olde Loohuis, D. P. Orfanos, J. F. Pearson, T. L. Pitcher, B. Pütz, Y. Quidé, A. Ragothaman, F. M. Rashid, W. R. Reay, R. Redlich, C. S. Reinbold, J. Repple, G. Richard, B. C. Riedel, S. L. Risacher, C. S. Rocha, N. R. Mota, L. Salminen, A. Saremi, A. J. Saykin, F. Schlag, L. Schmaal, P. R. Schofield, R. Secolin, C. Y. Shapland, L. Shen, J. Shin, E. Shumskaya, I. E. Sønderby, E. Sprooten, K. E. Tansey, A. Teumer, A. Thalamuthu, D. Tordesillas-Gutiérrez, J. A. Turner, A. Uhlmann, C. L. Vallerga, D. van der Meer, M. M. J. van Donkelaar, L. van Eijk, T. G. M. van Erp, N. E. M. van Haren, D. van Rooij, M.-J. van Tol, J. H. Veldink, E. Verhoef, E. Walton, M. Wang, Y. Wang, J. M. Wardlaw, W. Wen, L. T. Westlye, C. D. Whelan, S. H. Witt, K. Wittfeld, C. Wolf, T. Wolfers, J. Q. Wu, C. L. Yasuda, D. Zaremba, Z. Zhang, M. P. Zwiers, E. Artiges, A. A. Assareh, R. Ayesa-Arriola, A. Belger, C. L. Brandt, G. G. Brown, S. Cichon, J. E. Curran, G. E. Davies, F. Degenhardt, M. F. Dennis, B. Dietsche, S. Djurovic, C. P. Doherty, R. Espiritu, D. Garijo, Y. Gil, P. A. Gowland, R. C. Green, A. N. Häusler, W. Heindel, B.-C. Ho, W. U. Hoffmann, F. Holsboer, G. Homuth, N. Hosten, C. R. Jack, M. Jang, A. Jansen, N. A. Kimbrel, K. Kolskår, S. Koops, A. Krug, K. O. Lim, J. J. Luykx, D. H. Mathalon, K. A. Mather, V. S. Mattay, S. Matthews, J. van Mayoral Son, S. C. McEwen, I. Melle, D. W. Morris, B. A. Mueller, M. Nauck, J. E. Nordvik, M. M. Nöthen, D. S. O’Leary, N. Opel, M.-L. P. Martinot, G. B. Pike, A. Preda, E. B. Quinlan, P. E. Rasser, V. Ratnakar, S. Reppermund, V. M. Steen, P. A. Tooney, F. R. Torres, D. J. Veltman, J. T. Voyvodic, R. Whelan, T. White, H. Yamamori, H. H. H. Adams, J. C. Bis, S. Debette, C. Decarli, M. Fornage, V. Gudnason, E. Hofer, M. A. Ikram, L. Launer, W. T. Longstreth, O. L. Lopez, B. Mazoyer, T. H. Mosley, G. V. Roshchupkin, C. L. Satizabal, R. Schmidt, S. Seshadri, Q. Yang, M. K. M. Alvim, D. Ames, T. J. Anderson, O. A. Andreassen, A. Arias-Vasquez, M. E. Bastin, B. T. Baune, J. C. Beckham, J. Blangero, D. I. Boomsma, H. Brodaty, H. G. Brunner, R. L. Buckner, J. K. Buitelaar, J. R. Bustillo, W. Cahn, M. J. Cairns, V. Calhoun, V. J. Carr, X. Caseras, S. Caspers, G. L. Cavalleri, F. Cendes, A. Corvin, B. Crespo-Facorro, J. C. Dalrymple-Alford, U. Dannlowski, E. J. C. de Geus, I. J. Deary, N. Delanty, C. Depondt, S. Desrivières, G. Donohoe, T. Espeseth, G. Fernández, S. E. Fisher, H. Flor, A. J. Forstner, C. Francks, B. Franke, D. C. Glahn, R. L. Gollub, H. J. Grabe, O. Gruber, A. K. Håberg, A. R. Hariri, C. A. Hartman, R. Hashimoto, A. Heinz, F. A. Henskens, M. H. J. Hillegers, P. J. Hoekstra, A. J. Holmes, L. E. Hong, W. D. Hopkins, H. E. Hulshoff Pol, T. L. Jernigan, E. G. Jönsson, R. S. Kahn, M. A. Kennedy, T. T. J. Kircher, P. Kochunov, J. B. J. Kwok, S. Le Hellard, C. M. Loughland, N. G. Martin, J.-L. Martinot, C. McDonald, K. L. McMahon, A. Meyer-Lindenberg, P. T. Michie, R. A. Morey, B. Mowry, L. Nyberg, J. Oosterlaan, R. A. Ophoff, C. Pantelis, T. Paus, Z. Pausova, B. W. J. H. Penninx, T. J. C. Polderman, D. Posthuma, M. Rietschel, J. L. Roffman, L. M. Rowland, P. S. Sachdev, P. G. Sämann, U. Schall, G. Schumann, R. J. Scott, K. Sim, S. M. Sisodiya, J. W. Smoller, I. E. Sommer, B. St Pourcain, D. J. Stein, A. W. Toga, J. N. Trollor, N. J. A. van der Wee, D. van ‘t Ent, H. Völzke, H. Walter, B. Weber, D. R. Weinberger, M. J. Wright, J. Zhou, J. L. Stein, P. M. Thompson, S. E. Medland, The genetic architecture of the human cerebral cortex. Science 367, eaay6690 (2020).

34. J. J. Lee, M. McGue, W. G. Iacono, A. M. Michael, C. F. Chabris, The causal influence of brain size on human intelligence: Evidence from within-family phenotypic associations and GWAS modeling. Intelligence 75, 48–58 (2019).

35. S. Cheng, C. Wu, X. Qi, L. Liu, M. Ma, L. Zhang, B. Cheng, C. Liang, P. Li, O. P. Kafle, Y. Wen, F. Zhang, A Large-Scale Genetic Correlation Scan Between Intelligence and Brain Imaging Phenotypes. Cereb Cortex 30, 4197–4203 (2020).

36. J. Feng, C. Chen, Y. Cai, Z. Ye, K. Feng, J. Liu, L. Zhang, Q. Yang, A. Li, J. Sheng, B. Zhu, Z. Yu, C. Chen, Q. Dong, G. Xue, Partitioning heritability analyses unveil the genetic architecture of human brain multidimensional functional connectivity patterns. Hum Brain Mapp 41, 3305–3317 (2020).

37. T. Ge, C.-Y. Chen, A. E. Doyle, R. Vettermann, L. J. Tuominen, D. J. Holt, M. R. Sabuncu, J. W. Smoller, The Shared Genetic Basis of Educational Attainment and Cerebral Cortical Morphology. Cereb Cortex 29, 3471–3481 (2019).

38. P. R. Jansen, R. L. Muetzel, T. J. C. Polderman, V. W. Jaddoe, F. C. Verhulst, A. van der Lugt, H. Tiemeier, D. Posthuma, T. White, Polygenic Scores for Neuropsychiatric Traits and White Matter Microstructure in the Pediatric Population. Biol Psychiatry Cogn Neurosci Neuroimaging 4, 243–250 (2019).

39. M. Knol, A. Heshmatollah, L. Cremers, M. Kamran Ikram, A. Uitterlinden, C. van Dujin, W. Niessen, M. Vernooij, M. Afran Ikram, H. H. H. Adams, Genetic variation underlying cognition and its relation with neurological outcomes and brain imaging. Aging 11, 1440–1456 (2019).

40. Loughnan, R. J. et al, https://www.biorxiv.org/content/10.1101/637512v3 (2019).

41. M. L. Elliott, D. W. Belsky, K. Anderson, D. L. Corcoran, T. Ge, A. Knodt, J. A. Prinz, K. Sugden, B. Williams, D. Ireland, R. Poulton, A. Caspi, A. Holmes, T. Moffitt, A. R. Hariri, A Polygenic Score for Higher Educational Attainment is Associated with Larger Brains. Cereb Cortex 29, 3496–3504 (2019).

42. T. A. Lett, B. O. Vogel, S. Ripke, C. Wackerhagen, S. Erk, S. Awasthi, V. Trubetskoy, E. J. Brandl, S. Mohnke, I. M. Veer, M. M. Nöthen, M. Rietschel, F. Degenhardt, N. Romanczuk-Seiferth, S. H. Witt, T. Banaschewski, A. L. W. Bokde, C. Büchel, E. B. Quinlan, S. Desrivières, H. Flor, V. Frouin, H. Garavan, P. Gowland, B. Ittermann, J.-L. Martinot, M.-L. P. Martinot, F. Nees, D. Papadopoulos-Orfanos, T. Paus, L. Poustka, J. H. Fröhner, M. N. Smolka, R. Whelan, G. Schumann, H. Tost, A. Meyer-Lindenberg, A. Heinz, H. Walter, Cortical Surfaces Mediate the Relationship Between Polygenic Scores for Intelligence and General Intelligence. Cereb Cortex 30, 2707–2718 (2020).

43. B. L. Mitchell, G. Cuéllar-Partida, K. L. Grasby, A. I. Campos, L. T. Strike, L.-D. Hwang, A. Okbay, P. M. Thompson, S. E. Medland, N. G. Martin, M. J. Wright, M. E. Rentería, Educational attainment polygenic scores are associated with cortical total surface area and regions important for language and memory. NeuroImage 212, 116691 (2020).

44. E. Genç, C. Fraenz, C. Schlüter, P. Friedrich, M. C. Voelkle, R. Hossiep, O. Güntürkün, The Neural Architecture of General Knowledge. Eur J Pers 33, 589–605 (2019).

45. E. Genç, C. Fraenz, C. Schlüter, P. Friedrich, R. Hossiep, M. C. Voelkle, J. M. Ling, O. Güntürkün, R. E. Jung, Diffusion markers of dendritic density and arborization in gray matter predict differences in intelligence. Nat Commun 9, 1905 (2018).

46. E. Genç, C. Schlüter, C. Fraenz, L. Arning, D. Metzen, H. P. Nguyen, M. C. Voelkle, F. Streit, O. Güntürkün, R. Kumsta, S. Ocklenburg, Polygenic Scores for Cognitive Abilities and Their Association with Different Aspects of General Intelligence-A Deep Phenotyping Approach. Mol Neurobiol 58, 4145–4156 (2021).

47. A. Beauducel, B. Brocke, D. Liepmann, Perspectives on fluid and crystallized intelligence: facets for verbal, numerical, and figural intelligence. Personality and Individual Differences 30, 977–994 (2001).

48. D. Liepmann, A. Beauducel, B. Brocke, R. Amthauer, Intelligenz-Struktur-Test 2000 R (IST 2000 R). Manual (2. erweiterte und überarbeitete Aufl.) (Hogrefe, Göttingen, 2007).

49. L. A. Erdodi, C. A. Abeare, J. D. Lichtenstein, B. T. Tyson, B. Kucharski, B. G. Zuccato, R. M. Roth, Wechsler Adult Intelligence Scale-Fourth Edition (WAIS-IV) processing speed scores as measures of noncredible responding: The third generation of embedded performance validity indicators. Psychol Assess 29, 148–157 (2017).

50. A. M. Dale, B. Fischl, M. I. Sereno, Cortical surface-based analysis. I. Segmentation and surface reconstruction. NeuroImage 9, 179–194 (1999).

51. B. Fischl, M. I. Sereno, A. M. Dale, Cortical surface-based analysis. II: Inflation, flattening, and a surface-based coordinate system. NeuroImage 9, 195–207 (1999).

52. M. F. Glasser, T. S. Coalson, E. C. Robinson, C. D. Hacker, J. Harwell, E. Yacoub, K. Ugurbil, J. Andersson, C. F. Beckmann, M. Jenkinson, S. M. Smith, D. C. van Essen, A multi-modal parcellation of human cerebral cortex. Nature 536, 171–178 (2016).

53. B. Fischl, A. van der Kouwe, C. Destrieux, E. Halgren, F. Ségonne, D. H. Salat, E. Busa, L. J. Seidman, J. Goldstein, D. Kennedy, V. Caviness, N. Makris, B. Rosen, A. M. Dale, Automatically parcellating the human cerebral cortex. Cereb Cortex 14, 11–22 (2004).

54. T. E. J. Behrens, M. W. Woolrich, M. Jenkinson, H. Johansen-Berg, R. G. Nunes, S. Clare, P. M. Matthews, J. M. Brady, S. M. Smith, Characterization and propagation of uncertainty in diffusion-weighted MR imaging. Magn Reson Med 50, 1077–1088 (2003).

55. T. E. J. Behrens, H. J. Berg, S. Jbabdi, M. F. S. Rushworth, M. W. Woolrich, Probabilistic diffusion tractography with multiple fibre orientations: What can we gain? NeuroImage 34, 144–155 (2007).

56. M. Rubinov, O. Sporns, Complex network measures of brain connectivity: uses and interpretations. NeuroImage 52, 1059–1069 (2010).

57. M. Ivković, A. Kuceyeski, A. Raj, Statistics of weighted brain networks reveal hierarchical organization and Gaussian degree distribution. PloS one 7, e35029 (2012).

58. O. Sporns, D. R. Chialvo, M. Kaiser, C. C. Hilgetag, Organization, development and function of complex brain networks. Trends Cogn Sci 8, 418–425 (2004).

59. E. W. Dijkstra, A Note on Two Problems in Connexion with Graphs. Numerische Mathematik 1, 269–271 (1959).

60. A. Satorra, P. M. Bentler, in Latent variables analysis: Applications for developmental research, A. von Eye, C. C. Clogg, Eds. (Sage Publications, Thousand Oaks, CA, 1994), pp. 399–419.

61. S. Serang, R. Jacobucci, Exploratory Mediation Analysis of Dichotomous Outcomes via Regularization. Multivariate Behav Res 55, 69–86 (2020).

62. S. Serang, R. Jacobucci, K. C. Brimhall, K. J. Grimm, Exploratory Mediation Analysis via Regularization. Struct Equ Modeling 24, 733–744 (2017).

63. D. Góngora, M. Vega-Hernández, M. Jahanshahi, P. A. Valdés-Sosa, M. L. Bringas-Vega, Crystallized and fluid intelligence are predicted by microstructure of specific white-matter tracts. Hum Brain Mapp 41, 906–916 (2020).

64. B. A. Ammerman, S. Serang, R. Jacobucci, T. A. Burke, L. B. Alloy, M. S. McCloskey, Exploratory analysis of mediators of the relationship between childhood maltreatment and suicidal behavior. J Adolesc 69, 103–112 (2018).

65. H. Zou, T. Hastie, Regularization and variable selection via the elastic net. J R Statist Soc B 64, 301–320 (2005).

66. J. Friedman, T. Hastie, R. Tibshirani, Regularization Paths for Generalized Linear Models via Coordinate Descent. J Stat Soft 33(2010).

67. U. Basten, K. Hilger, C. J. Fiebach, Where smart brains are different: A quantitative meta-analysis of functional and structural brain imaging studies on intelligence. Intelligence 51, 10–27 (2015).

68. B. Pakkenberg, H. J. Gundersen, Neocortical neuron number in humans: effect of sex and age. J Comp Neurol 384, 312–320 (1997).

69. G. Leuba, R. Kraftsik, Changes in volume, surface estimate, three-dimensional shape and total number of neurons of the human primary visual cortex from midgestation until old age. Anat Embryol 190, 351–366 (1994).

70. J. Karbowski, Global and regional brain metabolic scaling and its functional consequences. BMC Biol 5, 18 (2007).

71. Z. Fatahi, A. Ghorbani, M. Ismail Zibaii, A. Haghparast, Neural synchronization between the anterior cingulate and orbitofrontal cortices during effort-based decision making. Neurobiol Learn Mem 175, 107320 (2020).

72. J. D. Wallis, S. W. Kennerley, Contrasting reward signals in the orbitofrontal cortex and anterior cingulate cortex. Annals of the New York Academy of Sciences 1239, 33–42 (2011).

73. S. Ramsden, F. M. Richardson, G. Josse, M. S. C. Thomas, C. Ellis, C. Shakeshaft, M. L. Seghier, C. J. Price, Verbal and non-verbal intelligence changes in the teenage brain. Nature 479, 113–116 (2011).

74. H. J. Syväoja, A. Kankaanpää, L. Joensuu, J. Kallio, H. Hakonen, C. H. Hillman, T. H. Tammelin, The Longitudinal Associations of Fitness and Motor Skills with Academic Achievement. Med Sci Sports Exerc 51, 2050–2057 (2019).

75. A. Trecroci, M. Duca, L. Cavaggioni, A. Rossi, R. Scurati, S. Longo, G. Merati, G. Alberti, D. Formenti, Relationship between Cognitive Functions and Sport-Specific Physical Performance in Youth Volleyball Players. Brain Sci 11(2021).

76. E. Santarnecchi, A. Emmendorfer, A. Pascual-Leone, Dissecting the parieto-frontal correlates of fluid intelligence: A comprehensive ALE meta-analysis study. Intelligence 63, 9–28 (2017).

77. J. D. Power, S. E. Petersen, Control-related systems in the human brain. Curr Opin Neurobiol 23, 223–228 (2013).

78. H. J. Park, K. Friston, Structural and functional brain networks: from connections to cognition. Science 342, 1238411 (2013).

79. A. Abdellaoui, K. J. H. Verweij, Dissecting polygenic signals from genome-wide association studies on human behaviour. Nat Hum Behav 5, 686–694 (2021).

80. A. Kong, G. Thorleifsson, M. L. Frigge, B. J. Vilhjalmsson, A. I. Young, T. E. Thorgeirsson, S. Benonisdottir, A. Oddsson, B. V. Halldorsson, G. Masson, D. F. Gudbjartsson, A. Helgason, G. Bjornsdottir, U. Thorsteinsdottir, K. Stefansson, The nature of nurture: Effects of parental genotypes. Science 359, 424–428 (2018).

