## Supplemental Material for "Structural architecture and brain network efficiency links polygenic scores to intelligence"

**Table S1. HCPMMP that are part of the P-FIT network**. This table shows all HCPMMP that overlap with BAs of the P-FIT network at least 80% in both hemispheres. The first column states the HCPMMP, the second column states the overlapping BAs belonging to the P-FIT network, in both hemispheres respectively. Brackets show the percentage of overlap.

| **HCPMMP** | **BA (%)** |
|---|---|
| 10d | LH_10 (87.8), LH_9 (8.7), LH_32 (3.5), RH_10 (99), RH_9 (1) |
| 10r | LH_32 (57.4), LH_10 (36.9), LH_24 (4.9), RH_10 (52.5), RH_32 (31.5) |
| 23d | LH_24 (72.5), LH_31 (14.3), RH_24 (79.9), RH_31 (11.6) |
| 24dv | LH_6 (55.7), LH_24 (44.3), RH_6 (70), RH_24 (23.3), RH_32 (3.1) |
| 31a | LH_31 (88.8), RH_31 (97.7) |
| 44 | LH_44 (89.9), LH_45 (2.4), RH_44 (62), RH_45 (33.5) |
| 45 | LH_45 (63.3), LH_47 (18.9), LH_44 (2.4), RH_45 (61.6), RH_47 (34.1) |
| 46 | LH_46 (66.2), LH_9 (32.2), LH_8 (1.6), RH_46 (66.8), RH_9 (30.6), RH_8 (2.1) |
| 6ma | LH_6 (87.4), RH_6 (95.9) |
| 6r | LH_6 (78.4), LH_44 (16.1), RH_6 (45.4), RH_44 (44.8) |
| 7PL | LH_7 (100), RH_7 (100) |
| 7Pm | LH_7 (99.8), LH_31 (0.2), RH_7 (92.2), RH_31 (7.8) |
| 8Ad | LH_8 (84.9), LH_6 (12.5), LH_9 (2.7), RH_8 (90.6), RH_9 (9.4) |
| 8Av | LH_8 (66.9), LH_9 (23.2), LH_6 (9.9), RH_8 (56.3), RH_9 (39.9), RH_6 (3.8) |
| 8BL | LH_8 (100), RH_8 (79.6), RH_9 (19.6) |
| 8BM | LH_8 (62.8), LH_32 (20.5), LH_9 (15.1), RH_8 (82.8), RH_9 (11.5), RH_32 (4.2) |
| 8C | LH_9 (42.1), LH_44 (34.1), LH_46 (19.7), RH_44 (38.3), RH_9 (34.8), RH_46 (26.1) |
| 9-46d | LH_10 (49.1), LH_9 (39.9), LH_46 (10.3), RH_9 (54.9), RH_10 (43), RH_46 (2.1) |
| 9a | LH_9 (94.5), LH_10 (5.5), RH_9 (79.6), RH_10 (20.4) |
| 9m | LH_9 (62.6), LH_32 (22.2), LH_10 (7.3), RH_9 (69.4), RH_32 (15.6), RH_10 (13.6) |
| 9p | LH_9 (86.4), LH_8 (13.6), RH_9 (100) |
| A1 | LH_42 (96.3), RH_42 (77) |
| DVT | LH_19 (87.3), LH_7 (11.4), LH_18 (1.3), RH_19 (69.3), RH_7 (29.6), RH_31 (0.7) |
| FFC | LH_37 (96.4), LH_19 (3.6), RH_37 (98.4), RH_19 (1.6) |
| FST | LH_39 (38.4), LH_37 (27.6), LH_19 (21.8), RH_37 (86), RH_19 (12.5) |
| IFJa | LH_44 (100), RH_44 (89.6), RH_45 (10.4) |
| IFJp | LH_6 (61.5), LH_44 (37.6), LH_9 (0.9), RH_44 (75.2), RH_6 (24.8) |
| IFSa | LH_46 (58.1), LH_45 (41.7), LH_47 (0.3), RH_46 (54.9), RH_45 (27.5), RH_47 (17.6) |
| IFSp | LH_44 (39.4), LH_45 (37), LH_46 (22.9), RH_45 (76.9), RH_46 (20.2), RH_44 (2.4) |
| IP0 | LH_19 (100), RH_19 (100) |
| IP1 | LH_7 (74.1), LH_19 (25.9), RH_7 (53.9), RH_19 (45.5) |
| IP2 | LH_7 (100), RH_7 (87.9), RH_40 (11.6) |
| IPS1 | LH_19 (91.5), LH_7 (8.5), RH_19 (95.3), RH_7 (4.7) |
| LIPd | LH_7 (100), RH_7 (100) |
| LIPv | LH_7 (100), RH_7 (100) |
| LO1 | LH_19 (100), RH_19 (100) |
| LO2 | LH_19 (100), RH_19 (96.6), RH_37 (3.4) |
| LO3 | LH_19 (98.6), LH_39 (1.4), RH_19 (100) |
| MIP | LH_7 (91.5), LH_19 (8.5), RH_7 (69.3), RH_19 (30.7) |
| MST | LH_39 (65.6), LH_19 (34.4), RH_19 (69.4), RH_39 (26.9), RH_37 (0.5) |
| MT | LH_19 (53.7), LH_39 (46.3), RH_19 (97.3), RH_39 (2.7) |

| | |
|---|---|
| PEF | LH_6 (100), RH_6 (99.3), RH_9 (0.8) |
| PF | LH_40 (84.6), LH_7 (15), RH_40 (94.3) |
| PFm | LH_7 (54.3), LH_40 (43.4), LH_39 (1.4), RH_40 (70.5), RH_7 (20.8), RH_39 (8.4) |
| PGi | LH_39 (62.8), LH_40 (37.2), RH_39 (90.1), RH_40 (9.7), RH_19 (0.2) |
| PGp | LH_19 (85.7), LH_39 (13.7), RH_19 (99.9), RH_39 (0.1) |
| PGs | LH_39 (52.4), LH_7 (27.8), LH_19 (16.6), RH_39 (45.7), RH_19 (27.9), RH_7 (17) |
| PHA2 | LH_20 (100), RH_20 (95.6) |
| PHA3 | LH_20 (74.3), LH_37 (25.7), RH_20 (77.1), RH_37 (22.1) |
| PH | LH_37 (98.4), LH_19 (1.6), RH_37 (99.9), RH_19 (0.1) |
| PIT | LH_19 (98.2), LH_37 (1.8), RH_19 (89.1), RH_37 (10.9) |
| POS2 | LH_7 (40.8), LH_31 (29.4), LH_19 (28.5), RH_31 (55.8), RH_7 (28.4), RH_19 (7.1) |
| SCEF | LH_6 (77.2), LH_8 (8.6), LH_32 (7), RH_6 (79.9), RH_8 (18.6) |
| SFL | LH_6 (72.8), LH_8 (27.2), RH_6 (55.1), RH_8 (44.9) |
| TE2a | LH_21 (75.8), LH_20 (23.5), RH_21 (71.3), RH_20 (22.6), RH_37 (4.7) |
| TE2p | LH_37 (60.8), LH_20 (22.5), LH_21 (15), RH_37 (87.1), RH_20 (10.5) |
| TF | LH_20 (88.4), LH_37 (11.6), RH_20 (74), RH_37 (26) |
| TPOJ2 | LH_39 (100), RH_39 (78.8), RH_37 (6.5) |
| TPOJ3 | LH_39 (96.9), LH_19 (3.1), RH_39 (66), RH_19 (34) |
| V1 | LH_18 (33.8), RH_18 (58.9), RH_19 (1.7) |
| V3A | LH_18 (97.6), LH_19 (2.4), RH_18 (95.6), RH_19 (4.4) |
| V3B | LH_19 (97.1), LH_18 (2.9), RH_18 (56.6), RH_19 (43.4) |
| V3CD | LH_19 (100), RH_19 (73.4), RH_18 (26.6) |
| V3 | LH_18 (60.2), LH_19 (39.8), RH_18 (55.5), RH_19 (44.5) |
| V4 | LH_19 (71.7), LH_18 (28.3), RH_19 (72.8), RH_18 (27.2) |
| V4t | LH_19 (100), RH_19 (97.6), RH_37 (2.4) |
| V6A | LH_19 (69.3), LH_18 (30.7), RH_19 (97), RH_7 (3) |
| V6 | LH_18 (58.4), LH_19 (40.9), RH_19 (87.2), RH_18 (12.3) |
| V7 | LH_19 (71.9), LH_18 (28.1), RH_19 (55.5), RH_18 (44.5) |
| V8 | LH_19 (74.5), LH_37 (22.9), RH_19 (83.8), RH_37 (14.8) |
| VIP | LH_7 (100), RH_7 (100) |
| VMV2 | LH_37 (50.3), LH_19 (29.9), LH_20 (16.6), RH_20 (53.4), RH_19 (24.7), RH_37 (20.3) |
| VMV3 | LH_37 (59.6), LH_19 (40.4), RH_37 (64), RH_19 (34.5) |
| VVC | LH_37 (88.4), LH_20 (11.3), LH_19 (0.3), RH_37 (92.6), RH_20 (6.5), RH_19 (0.9) |
| a24pr | LH_24 (100), RH_24 (100) |
| a32pr | LH_32 (66.4), LH_24 (33.6), RH_32 (63.5), RH_24 (35.4), RH_8 (1.1) |
| a9-46v | LH_10 (83.6), LH_46 (16.4), RH_10 (86.9), RH_46 (13.1) |
| d32 | LH_32 (86.9), LH_24 (6.6), RH_32 (73.8), RH_9 (21.5), RH_24 (4.2) |
| i6-8 | LH_6 (94.2), LH_8 (5.8), RH_6 (52.3), RH_8 (47.7) |
| p10p | LH_10 (100), RH_10 (99.1) |
| p24 | LH_24 (96.7), RH_24 (92.5) |
| p24pr | LH_24 (100), RH_24 (100) |
| p32 | LH_32 (63.4), LH_24 (36.4), RH_32 (65.9), RH_24 (31.6), RH_10 (2.5) |
| p32pr | LH_24 (56.6), LH_32 (35.7), LH_6 (3.5), RH_32 (49.4), RH_24 (34.3), RH_8 (10.3) |
| p47r | LH_46 (56), LH_47 (24.4), LH_10 (18.4), RH_47 (52.8), RH_10 (24), RH_46 (18.1) |
| p9-46v | LH_46 (92.7), LH_9 (7.3), RH_46 (99.8), RH_9 (0.2) |
| s32 | LH_24 (71.8), LH_32 (26.8), RH_32 (64.6), RH_24 (29) |
| s6-8 | LH_6 (56.2), LH_8 (43.8), RH_8 (59.1), RH_6 (40.9) |

**Table S2. Path a coefficients**. Coefficients from the a path (PGS - brain) of the mediation analysis after re-estimation with lavaan. Analyses were conducted with two different PGS (EA, GI). Results from the surface area (SA) analysis are depicted in green, results from the cortical thickness (CT) analysis in blue, results from the DWI analysis in red, results from the rsfMRI anaylsis in yellow.

| PGS-EA | | PGS-GI | | PGS-EA | | PGS-GI | | PGS-EA | | PGS-GI | | PGS-EA | | PGS-GI | |
|---|---|---|---|---|---|---|---|---|---|---|---|---|---|---|---|
| UV | a | UV | a | UV | a | UV | a | UV | a | UV | a | UV | a | UV | a |
| SA_V1 | 0.076 | SA_V1 | 0.055 | CT_V1 | -0.042 | CT_V1 | -0.039 | DWI_V1 | 0.112 | DWI_MST | 0.071 | rsfMRI_MST | 0.058 | rsfMRI_V2 | 0.053 |
| SA_MST | 0.100 | SA_V2 | 0.053 | CT_MT | -0.099 | CT_FEF | 0.048 | DWI_MST | 0.143 | DWI_4 | 0.083 | rsfMRI_V2 | 0.053 | rsfMRI_3b | 0.058 |
| SA_V6 | 0.064 | SA_4 | 0.079 | CT_PSL | -0.042 | CT_PEF | 0.045 | DWI_V6 | 0.109 | DWI_3b | 0.079 | rsfMRI_V3 | 0.049 | rsfMRI_PEF | -0.055 |
| SA_V2 | 0.086 | SA_3b | 0.072 | CT_STV | -0.050 | CT_V7 | -0.085 | DWI_V2 | 0.115 | DWI_FEF | 0.112 | rsfMRI_4 | 0.045 | rsfMRI_POS2 | -0.053 |
| SA_V3 | 0.077 | SA_FEF | 0.106 | CT_POS1 | -0.072 | CT_LO2 | 0.061 | DWI_V3 | 0.114 | DWI_PEF | 0.081 | rsfMRI_3b | 0.045 | rsfMRI_LO1 | 0.043 |
| SA_V4 | 0.089 | SA_PEF | 0.079 | CT_23d | 0.057 | CT_PIT | 0.041 | DWI_V4 | 0.113 | DWI_55b | 0.096 | rsfMRI_A1 | 0.079 | rsfMRI_5m | 0.059 |
| SA_4 | 0.094 | SA_55b | 0.082 | CT_d23ab | -0.052 | CT_PSL | -0.056 | DWI_V8 | 0.113 | DWI_IPS1 | 0.046 | rsfMRI_POS1 | 0.052 | rsfMRI_5L | 0.054 |
| SA_3b | 0.083 | SA_IPS1 | 0.050 | CT_5L | 0.044 | CT_SCEF | -0.066 | DWI_4 | 0.133 | DWI_FFC | 0.080 | rsfMRI_5m | 0.090 | rsfMRI_1 | 0.061 |
| SA_FEF | 0.083 | SA_SFL | 0.084 | CT_24dd | 0.046 | CT_MIP | 0.057 | DWI_3b | 0.126 | DWI_MT | 0.059 | rsfMRI_5mv | 0.061 | rsfMRI_3a | 0.064 |
| SA_PEF | 0.101 | SA_PCV | 0.060 | CT_7Am | 0.048 | CT_6v | 0.047 | DWI_FEF | 0.158 | DWI_A1 | 0.068 | rsfMRI_24dd | 0.073 | rsfMRI_33pr | 0.048 |
| SA_55b | 0.058 | SA_STV | 0.062 | CT_7PL | -0.061 | CT_a24pr | -0.077 | DWI_PEF | 0.134 | DWI_SFL | 0.082 | rsfMRI_24dv | 0.047 | rsfMRI_a24pr | -0.053 |
| SA_V3A | 0.057 | SA_7m | 0.067 | CT_a24 | 0.087 | CT_47m | -0.050 | DWI_55b | 0.156 | DWI_PCV | 0.060 | rsfMRI_2 | 0.054 | rsfMRI_a24 | 0.077 |
| SA_RSC | 0.102 | SA_POS1 | 0.068 | CT_d32 | 0.069 | CT_10d | -0.068 | DWI_V3A | 0.124 | DWI_STV | 0.070 | rsfMRI_3a | 0.055 | rsfMRI_s68 | -0.057 |
| SA_V7 | 0.093 | SA_23d | 0.053 | CT_8BM | 0.068 | CT_45 | -0.060 | DWI_RSC | 0.128 | DWI_7Pm | 0.054 | rsfMRI_p24pr | 0.058 | rsfMRI_52 | 0.062 |
| SA_IPS1 | 0.064 | SA_v23ab | 0.069 | CT_p32 | 0.063 | CT_a47r | -0.051 | DWI_POS2 | 0.099 | DWI_23d | 0.043 | rsfMRI_33pr | 0.066 | rsfMRI_ProS | 0.068 |
| SA_FFC | 0.056 | SA_31pv | 0.044 | CT_8Av | 0.041 | CT_IFJa | -0.047 | DWI_V7 | 0.135 | DWI_31pv | 0.060 | rsfMRI_a24pr | 0.086 | rsfMRI_PHA1 | 0.053 |
| SA_V3B | 0.066 | SA_24dd | 0.077 | CT_9m | 0.060 | CT_p946v | -0.055 | DWI_IPS1 | 0.132 | DWI_5m | 0.058 | rsfMRI_a24 | 0.062 | rsfMRI_TE2a | -0.055 |
| SA_LO1 | 0.095 | SA_24dv | 0.066 | CT_44 | -0.058 | CT_10pp | -0.072 | DWI_FFC | 0.130 | DWI_5mv | 0.055 | rsfMRI_8BM | 0.054 | rsfMRI_PGs | -0.062 |
| SA_LO2 | 0.081 | SA_SCEF | 0.098 | CT_a946v | 0.041 | CT_11l | -0.074 | DWI_V3B | 0.103 | DWI_23c | 0.059 | rsfMRI_p32 | 0.068 | rsfMRI_VMV1 | 0.082 |
| SA_PIT | 0.068 | SA_6ma | 0.079 | CT_9a | 0.079 | CT_13l | -0.068 | DWI_LO1 | 0.136 | DWI_5L | 0.068 | rsfMRI_47m | 0.099 | rsfMRI_PHA2 | 0.045 |
| SA_MT | 0.086 | SA_7PL | 0.063 | CT_a10p | 0.059 | CT_43 | 0.043 | DWI_LO2 | 0.108 | DWI_24dd | 0.061 | rsfMRI_9p | 0.074 | rsfMRI_LO3 | 0.042 |
| SA_SFL | 0.079 | SA_LIPv | 0.062 | CT_6a | 0.054 | CT_OP1 | -0.057 | DWI_PIT | 0.105 | DWI_24dv | 0.064 | rsfMRI_10d | 0.060 | rsfMRI_VMV2 | 0.054 |
| SA_STV | 0.088 | SA_VIP | 0.045 | CT_RI | -0.060 | CT_OP23 | -0.058 | DWI_MT | 0.141 | DWI_7AL | 0.060 | rsfMRI_44 | 0.046 | rsfMRI_MBelt | 0.073 |
| SA_POS1 | 0.068 | SA_MIP | 0.083 | CT_AVI | 0.074 | CT_RI | -0.048 | DWI_A1 | 0.128 | DWI_SCEF | 0.082 | rsfMRI_45 | 0.059 | rsfMRI_LBelt | 0.060 |
| SA_23d | 0.083 | SA_1 | 0.044 | CT_FOP1 | 0.115 | CT_MI | -0.076 | DWI_PSL | 0.105 | DWI_6ma | 0.091 | rsfMRI_47l | 0.087 | rsfMRI_A4 | 0.047 |
| SA_v23ab | 0.084 | SA_3a | 0.073 | CT_FOP3 | 0.069 | CT_Pir | -0.071 | DWI_SFL | 0.116 | DWI_7PC | 0.064 | rsfMRI_a47r | 0.053 | | |
| SA_d23ab | 0.086 | SA_6d | 0.095 | CT_PreS | 0.069 | CT_FOP1 | 0.096 | DWI_PCV | 0.092 | DWI_LIPv | 0.075 | rsfMRI_9a | 0.055 | | |
| SA_31pv | 0.055 | SA_6mp | 0.080 | CT_ProS | -0.070 | CT_FOP3 | 0.041 | DWI_STV | 0.138 | DWI_VIP | 0.071 | rsfMRI_10v | 0.065 | | |
| SA_23c | 0.046 | SA_p32pr | 0.069 | CT_PeEc | 0.059 | CT_FOP2 | 0.064 | DWI_7Pm | 0.088 | DWI_MIP | 0.063 | rsfMRI_a10p | 0.065 | | |
| SA_6ma | 0.093 | SA_a24 | 0.056 | CT_TGd | 0.075 | CT_AIP | 0.046 | DWI_7m | 0.110 | DWI_1 | 0.079 | rsfMRI_10pp | 0.116 | | |
| SA_7PL | 0.080 | SA_d32 | 0.062 | CT_PHT | 0.066 | CT_PreS | 0.064 | DWI_POS1 | 0.115 | DWI_2 | 0.074 | rsfMRI_13l | 0.070 | | |
| SA_LIPv | 0.081 | SA_8BM | 0.059 | CT_PGp | -0.071 | CT_H | 0.056 | DWI_23d | 0.140 | DWI_3a | 0.086 | rsfMRI_47s | 0.071 | | |
| SA_VIP | 0.117 | SA_p32 | 0.054 | CT_IP1 | -0.077 | CT_ProS | -0.064 | DWI_v23ab | 0.122 | DWI_6d | 0.097 | rsfMRI_6a | 0.047 | | |
| SA_MIP | 0.076 | SA_47m | 0.085 | CT_31pd | -0.047 | CT_STGa | 0.053 | DWI_d23ab | 0.135 | DWI_6mp | 0.072 | rsfMRI_43 | 0.064 | | |
| SA_1 | 0.059 | SA_8Ad | 0.064 | CT_25 | 0.075 | CT_PHA1 | 0.065 | DWI_31pv | 0.144 | DWI_6v | 0.060 | rsfMRI_OP4 | 0.064 | | |
| SA_3a | 0.114 | SA_8BL | 0.081 | CT_FOP5 | 0.078 | CT_PHA3 | 0.049 | DWI_5m | 0.106 | DWI_p24pr | 0.054 | rsfMRI_OP1 | 0.045 | | |
| SA_6d | 0.116 | SA_9p | 0.073 | CT_p10p | 0.043 | CT_STSdp | 0.051 | DWI_5mv | 0.125 | DWI_33pr | 0.043 | rsfMRI_OP23 | 0.087 | | |
| SA_6mp | 0.066 | SA_10d | 0.046 | CT_a32pr | 0.091 | CT_PH | 0.086 | DWI_23c | 0.112 | DWI_a24pr | 0.065 | rsfMRI_52 | 0.113 | | |
| SA_6v | 0.062 | SA_a47r | 0.081 | CT_p24 | 0.067 | CT_TPOJ3 | -0.055 | DWI_5L | 0.122 | DWI_p32pr | 0.062 | rsfMRI_RI | 0.061 | | |
| SA_a24 | 0.062 | SA_6r | 0.083 | | | CT_IP1 | -0.054 | DWI_24dd | 0.117 | DWI_8BM | 0.055 | rsfMRI_PFcm | 0.072 | | |
| SA_d32 | 0.049 | SA_IFJa | 0.051 | | | CT_PGi | -0.052 | DWI_24dv | 0.108 | DWI_p32 | 0.051 | rsfMRI_PoI2 | 0.072 | | |
| SA_8BM | 0.050 | SA_IFJp | 0.068 | | | CT_FST | 0.060 | DWI_7AL | 0.088 | DWI_47m | 0.086 | rsfMRI_TA2 | 0.059 | | |
| SA_p32 | 0.057 | SA_46 | 0.041 | | | CT_VMV2 | 0.049 | DWI_SCEF | 0.105 | DWI_8Av | 0.100 | rsfMRI_Pir | 0.057 | | |
| SA_10r | 0.047 | SA_a946v | 0.092 | | | CT_25 | -0.054 | DWI_6ma | 0.140 | DWI_8Ad | 0.104 | rsfMRI_AAIC | 0.058 | | |
| SA_47m | 0.086 | SA_946d | 0.053 | | | CT_s32 | -0.065 | DWI_7Am | 0.085 | DWI_8BL | 0.084 | rsfMRI_FOP3 | 0.054 | | |
| SA_8Ad | 0.064 | SA_9a | 0.061 | | | CT_pOFC | -0.082 | DWI_7PL | 0.116 | DWI_9p | 0.096 | rsfMRI_PFt | 0.066 | | |
| SA_9m | 0.044 | SA_a10p | 0.062 | | | CT_TE1m | -0.062 | DWI_7PC | 0.098 | DWI_8C | 0.078 | rsfMRI_AIP | 0.065 | | |
| SA_8BL | 0.075 | SA_11l | 0.054 | | | CT_V2 | -0.003 | DWI_LIPv | 0.124 | DWI_44 | 0.065 | rsfMRI_ProS | 0.050 | | |
| SA_9p | 0.103 | SA_13l | 0.132 | | | | | DWI_VIP | 0.133 | DWI_45 | 0.054 | rsfMRI_PBelt | 0.047 | | |
| SA_10d | 0.059 | SA_47s | 0.076 | | | | | DWI_MIP | 0.135 | DWI_47l | 0.057 | rsfMRI_PHA1 | 0.060 | | |
| SA_44 | 0.051 | SA_6a | 0.104 | | | | | DWI_1 | 0.125 | DWI_a47r | 0.074 | rsfMRI_TGd | 0.059 | | |
| SA_a47r | 0.081 | SA_i68 | 0.088 | | | | | DWI_2 | 0.129 | DWI_6r | 0.066 | rsfMRI_TE1a | 0.057 | | |
| SA_6r | 0.067 | SA_s68 | 0.084 | | | | | DWI_3a | 0.134 | DWI_IFJa | 0.075 | rsfMRI_TPOJ1 | 0.058 | | |
| SA_IFJp | 0.062 | SA_43 | 0.072 | | | | | DWI_6d | 0.147 | DWI_IFJp | 0.077 | rsfMRI_DVT | 0.038 | | |
| SA_IFSa | -0.073 | SA_OP1 | 0.057 | | | | | DWI_6mp | 0.117 | DWI_IFSp | 0.074 | rsfMRI_PGp | 0.053 | | |
| SA_46 | 0.053 | SA_PoI2 | 0.043 | | | | | DWI_6v | 0.134 | DWI_IFSa | 0.078 | rsfMRI_V6A | 0.046 | | |
| SA_946d | 0.071 | SA_TA2 | 0.060 | | | | | DWI_p24pr | 0.118 | DWI_p946v | 0.092 | rsfMRI_FST | 0.068 | | |
| SA_9a | 0.119 | SA_AAIC | 0.060 | | | | | DWI_33pr | 0.127 | DWI_46 | 0.103 | rsfMRI_25 | 0.071 | | |
| SA_10v | 0.057 | SA_FOP1 | 0.057 | | | | | DWI_a24pr | 0.127 | DWI_a946v | 0.102 | rsfMRI_PoI1 | 0.081 | | |
| SA_a10p | 0.071 | SA_FOP3 | 0.079 | | | | | DWI_p32pr | 0.122 | DWI_946d | 0.106 | rsfMRI_Ig | 0.067 | | |
| SA_10pp | 0.107 | SA_FOP2 | 0.112 | | | | | DWI_a24 | 0.108 | DWI_9a | 0.072 | rsfMRI_MBelt | 0.081 | | |
| SA_11l | 0.100 | SA_EC | 0.056 | | | | | DWI_d32 | 0.110 | DWI_a10p | 0.066 | rsfMRI_LBelt | 0.076 | | |
| SA_13l | 0.101 | SA_PeEc | 0.047 | | | | | DWI_8BM | 0.103 | DWI_11l | 0.072 | rsfMRI_A4 | 0.062 | | |
| SA_OFC | 0.091 | SA_STSdp | 0.059 | | | | | DWI_p32 | 0.120 | DWI_13l | 0.072 | rsfMRI_PI | 0.060 | | |
| SA_47s | 0.061 | SA_STSvp | 0.058 | | | | | DWI_10r | 0.135 | DWI_47s | 0.098 | rsfMRI_cauda | 0.071 | | |
| SA_6a | 0.072 | SA_TE1p | 0.113 | | | | | DWI_47m | 0.136 | DWI_LIPd | 0.074 | rsfMRI_hippo | 0.060 | | |
| SA_i68 | 0.064 | SA_TF | 0.045 | | | | | DWI_8Av | 0.134 | DWI_6a | 0.094 | rsfMRI_puta | 0.078 | | |
| SA_s68 | 0.041 | SA_TE2p | 0.084 | | | | | DWI_8Ad | 0.139 | DWI_i68 | 0.096 | rsfMRI_thala | 0.049 | | |
| SA_OP1 | 0.070 | SA_PHT | 0.090 | | | | | DWI_9m | 0.113 | DWI_s68 | 0.116 | | | | |
| SA_OP23 | 0.056 | SA_PH | 0.084 | | | | | DWI_8BL | 0.114 | DWI_43 | 0.082 | | | | |
| SA_52 | 0.081 | SA_PGp | 0.047 | | | | | DWI_9p | 0.125 | DWI_OP4 | 0.087 | | | | |
| SA_TA2 | 0.076 | SA_IP1 | 0.084 | | | | | DWI_10d | 0.091 | DWI_OP1 | 0.096 | | | | |
| SA_FOP4 | 0.057 | SA_PFm | 0.042 | | | | | DWI_8C | 0.116 | DWI_OP23 | 0.073 | | | | |
| SA_Pir | 0.052 | SA_VMV1 | 0.072 | | | | | DWI_44 | 0.136 | DWI_52 | 0.058 | | | | |
| SA_FOP3 | 0.064 | SA_PHA2 | 0.051 | | | | | DWI_45 | 0.111 | DWI_RI | 0.066 | | | | |
| SA_FOP2 | 0.051 | SA_FST | 0.049 | | | | | DWI_47l | 0.115 | DWI_PFcm | 0.072 | | | | |
| SA_PFt | 0.110 | SA_25 | 0.056 | | | | | DWI_a47r | 0.116 | DWI_PoI2 | 0.057 | | | | |
| SA_AIP | 0.071 | SA_s32 | 0.053 | | | | | DWI_6r | 0.124 | DWI_TA2 | 0.054 | | | | |
| SA_PreS | 0.053 | SA_pOFC | 0.060 | | | | | DWI_IFJa | 0.111 | DWI_FOP4 | 0.069 | | | | |
| SA_H | 0.046 | SA_PoI1 | 0.052 | | | | | DWI_IFJp | 0.115 | DWI_MI | 0.071 | | | | |
| SA_ProS | 0.078 | SA_Ig | 0.080 | | | | | DWI_IFSp | 0.103 | DWI_Pir | 0.054 | | | | |
| SA_PeEc | 0.047 | SA_p10p | 0.067 | | | | | DWI_IFSa | 0.082 | DWI_AVI | 0.061 | | | | |
| SA_STGa | 0.042 | SA_p47r | 0.085 | | | | | DWI_p946v | 0.111 | DWI_AAIC | 0.049 | | | | |
| SA_PHA1 | 0.069 | SA_MBelt | 0.072 | | | | | DWI_46 | 0.135 | DWI_FOP1 | 0.069 | | | | |
| SA_STSvp | 0.060 | SA_TE1m | 0.073 | | | | | DWI_a946v | 0.129 | DWI_FOP3 | 0.069 | | | | |
| SA_TE1p | 0.096 | SA_PI | 0.079 | | | | | DWI_946d | 0.129 | DWI_FOP2 | 0.067 | | | | |
| SA_TF | 0.045 | SA_a32pr | 0.045 | | | | | DWI_9a | 0.114 | DWI_PFt | 0.066 | | | | |
| SA_TE2p | 0.071 | | | | | | | DWI_10v | 0.127 | DWI_AIP | 0.078 | | | | |

| | | | | | |
|---|---|---|---|---|---|
| SA_PHT | 0.131 | DWI_a10p | 0.124 | DWI_EC | 0.059 |
| SA_PH | 0.099 | DWI_10pp | 0.114 | DWI_H | 0.057 |
| SA_TPOJ1 | 0.074 | DWI_11l | 0.152 | DWI_PeEc | 0.055 |
| SA_TPOJ2 | 0.080 | DWI_13l | 0.148 | DWI_STGa | 0.063 |
| SA_DVT | 0.048 | DWI_OFC | 0.124 | DWI_PBelt | 0.071 |
| SA_IP2 | 0.069 | DWI_47s | 0.139 | DWI_A5 | 0.076 |
| SA_IP1 | 0.070 | DWI_LIPd | 0.126 | DWI_PHA3 | 0.074 |
| SA_PFop | 0.051 | DWI_6a | 0.140 | DWI_STSda | 0.075 |
| SA_PFm | 0.058 | DWI_i68 | 0.135 | DWI_STSdp | 0.086 |
| SA_VMV1 | 0.110 | DWI_s68 | 0.133 | DWI_STSvp | 0.088 |
| SA_PHA2 | 0.049 | DWI_43 | 0.132 | DWI_TGd | 0.057 |
| SA_V4t | 0.089 | DWI_OP4 | 0.148 | DWI_TE1a | 0.066 |
| SA_FST | 0.116 | DWI_OP1 | 0.156 | DWI_TE1p | 0.104 |
| SA_V3CD | 0.093 | DWI_OP23 | 0.133 | DWI_TE2a | 0.089 |
| SA_LO3 | 0.099 | DWI_52 | 0.114 | DWI_TF | 0.091 |
| SA_VMV2 | 0.071 | DWI_RI | 0.109 | DWI_TE2p | 0.098 |
| SA_25 | 0.048 | DWI_PFcm | 0.117 | DWI_PHT | 0.100 |
| SA_s32 | 0.047 | DWI_PoI2 | 0.118 | DWI_PH | 0.094 |
| SA_pOFC | 0.066 | DWI_TA2 | 0.111 | DWI_TPOJ1 | 0.079 |
| SA_PoI1 | 0.070 | DWI_FOP4 | 0.135 | DWI_TPOJ2 | 0.086 |
| SA_MBelt | 0.096 | DWI_MI | 0.131 | DWI_TPOJ3 | 0.055 |
| SA_TE1m | 0.088 | DWI_Pir | 0.092 | DWI_PGp | 0.052 |
| SA_PI | 0.077 | DWI_AVI | 0.126 | DWI_IP2 | 0.073 |
| SA_a32pr | 0.059 | DWI_AAIC | 0.075 | DWI_IP1 | 0.073 |
| | | DWI_FOP1 | 0.128 | DWI_IP0 | 0.054 |
| | | DWI_FOP3 | 0.132 | DWI_PFop | 0.067 |
| | | DWI_FOP2 | 0.122 | DWI_PF | 0.059 |
| | | DWI_PFt | 0.123 | DWI_PFm | 0.075 |
| | | DWI_AIP | 0.135 | DWI_PGi | 0.077 |
| | | DWI_EC | 0.120 | DWI_PGs | 0.081 |
| | | DWI_PreS | 0.126 | DWI_V6A | 0.050 |
| | | DWI_H | 0.127 | DWI_PHA2 | 0.057 |
| | | DWI_ProS | 0.126 | DWI_V4t | 0.062 |
| | | DWI_PeEc | 0.126 | DWI_FST | 0.079 |
| | | DWI_STGa | 0.093 | DWI_31pd | 0.051 |
| | | DWI_PBelt | 0.123 | DWI_31a | 0.052 |
| | | DWI_A5 | 0.127 | DWI_VVC | 0.064 |
| | | DWI_PHA1 | 0.122 | DWI_s32 | 0.055 |
| | | DWI_PHA3 | 0.137 | DWI_PoI1 | 0.070 |
| | | DWI_STSda | 0.122 | DWI_Ig | 0.077 |
| | | DWI_STSdp | 0.132 | DWI_FOP5 | 0.056 |
| | | DWI_STSvp | 0.127 | DWI_p10p | 0.078 |
| | | DWI_TGd | 0.096 | DWI_p47r | 0.073 |
| | | DWI_TE1a | 0.109 | DWI_TGv | 0.062 |
| | | DWI_TE1p | 0.139 | DWI_MBelt | 0.069 |
| | | DWI_TE2a | 0.122 | DWI_LBelt | 0.064 |
| | | DWI_TF | 0.126 | DWI_A4 | 0.068 |
| | | DWI_TE2p | 0.130 | DWI_STSva | 0.077 |
| | | DWI_PHT | 0.164 | DWI_TE1m | 0.082 |
| | | DWI_PH | 0.143 | DWI_PI | 0.063 |
| | | DWI_TPOJ1 | 0.128 | DWI_cauda | 0.058 |
| | | DWI_TPOJ2 | 0.149 | DWI_hippo | 0.053 |
| | | DWI_TPOJ3 | 0.129 | DWI_palli | 0.063 |
| | | DWI_DVT | 0.122 | DWI_puta | 0.066 |
| | | DWI_PGp | 0.125 | DWI_thala | 0.059 |
| | | DWI_IP2 | 0.130 | DWI_ventdc | 0.060 |
| | | DWI_IP1 | 0.134 | | |
| | | DWI_IP0 | 0.122 | | |
| | | DWI_PFop | 0.109 | | |
| | | DWI_PF | 0.116 | | |
| | | DWI_PFm | 0.141 | | |
| | | DWI_PGi | 0.141 | | |
| | | DWI_PGs | 0.142 | | |
| | | DWI_V6A | 0.114 | | |
| | | DWI_VMV1 | 0.125 | | |
| | | DWI_VMV3 | 0.118 | | |
| | | DWI_PHA2 | 0.146 | | |
| | | DWI_V4t | 0.128 | | |
| | | DWI_FST | 0.150 | | |
| | | DWI_V3CD | 0.107 | | |
| | | DWI_LO3 | 0.132 | | |
| | | DWI_VMV2 | 0.114 | | |
| | | DWI_31pd | 0.118 | | |
| | | DWI_31a | 0.130 | | |
| | | DWI_VVC | 0.120 | | |
| | | DWI_25 | 0.101 | | |
| | | DWI_s32 | 0.122 | | |
| | | DWI_pOFC | 0.098 | | |
| | | DWI_PoI1 | 0.125 | | |
| | | DWI_Ig | 0.124 | | |
| | | DWI_FOP5 | 0.118 | | |
| | | DWI_p10p | 0.126 | | |
| | | DWI_p47r | 0.118 | | |
| | | DWI_TGv | 0.102 | | |
| | | DWI_MBelt | 0.135 | | |
| | | DWI_LBelt | 0.123 | | |
| | | DWI_A4 | 0.128 | | |
| | | DWI_STSva | 0.112 | | |
| | | DWI_TE1m | 0.132 | | |
| | | DWI_PI | 0.116 | | |
| | | DWI_a32pr | 0.119 | | |
| | | DWI_p24 | 0.101 | | |
| | | DWI_accumb | 0.103 | | |
| | | DWI_amy | 0.107 | | |
| | | DWI_cauda | 0.116 | | |
| | | DWI_hippo | 0.122 | | |
| | | DWI_palli | 0.132 | | |
| | | DWI_puta | 0.118 | | |
| | | DWI_thala | 0.118 | | |
| | | DWI_ventdc | 0.122 | | |

**Table S3. Path b coefficients**. Coefficients from the b path (brain - IST 2000 R score) of the mediation analysis after re-estimation with lavaan. Analyses were performed twice, controlling for one of the PGS (EA, GI). Results from the SA analysis are depicted in green, results from the CT analysis in blue and results from the DWI analysis in red. There were no brain areas whose rsfMRI showed any association with I-ST 200 R score.

| PGS-EA | | PGS-GI | | PGS-EA | | PGS-GI | | PGS-EA | | PGS-GI | |
|---|---|---|---|---|---|---|---|---|---|---|---|
| **UV** | **b** | **UV** | **b** | **UV** | **b** | **UV** | **b** | **UV** | **b** | **UV** | **b** |
| SA_4 | 0.078 | SA_MIP | 0.056 | CT_FOP1 | 0.122 | | | DWI_LO1 | -0.050 | DWI_45 | 0.338 |
| SA_5mv | -0.120 | SA_2 | -0.190 | CT_PreS | 0.120 | | | DWI_LO2 | -0.102 | DWI_Pir | -0.138 |
| SA_5L | 0.009 | SA_45 | 0.109 | CT_H | 0.086 | | | DWI_5mv | -0.251 | DWI_VMV1 | -0.110 |
| SA_7PC | -0.080 | SA_OFC | 0.089 | CT_TE2a | -0.104 | | | DWI_6ma | 0.153 | | |
| SA_MIP | 0.065 | SA_OP1 | 0.060 | CT_25 | -0.048 | | | DWI_44 | 0.068 | | |
| SA_1 | -0.227 | SA_STGa | 0.068 | CT_pOFC | -0.113 | | | DWI_45 | 0.164 | | |
| SA_2 | 0.037 | SA_PH | 0.060 | CT_PI | 0.100 | | | DWI_6r | 0.136 | | |
| SA_45 | 0.049 | SA_IP1 | 0.051 | | | | | DWI_47s | 0.234 | | |
| SA_6r | 0.055 | | | | | | | DWI_43 | 0.159 | | |
| SA_IFSa | 0.121 | | | | | | | DWI_Pir | -0.210 | | |
| SA_OFC | 0.064 | | | | | | | DWI_VMV1 | -0.157 | | |
| SA_OP1 | 0.059 | | | | | | | DWI_pOFC | -0.057 | | |
| SA_FOP3 | 0.035 | | | | | | | | | | |
| SA_EC | 0.094 | | | | | | | | | | |
| SA_ProS | -0.091 | | | | | | | | | | |
| SA_STGa | 0.078 | | | | | | | | | | |
| SA_PH | 0.068 | | | | | | | | | | |
| SA_IP1 | 0.043 | | | | | | | | | | |

**Table S4. Mediation coefficients**. Coefficients of the mediation analysis after re-estimation with lavaan. Tables show coefficients for a, b and ab. Analyses were performed two times, with three different PGS as IV (EA, GI). Results from the SA analysis are depicted in green, results from the CT analysis in blue and results from the DWI analysis in red. There are no areas whose rsfMRI connectivity mediated the effect of any of the PGS on I-S-T 2000 R score.

| PGS-EA | | | | PGS-GI | | | | PGS-EA | | | | PGS-GI | | | | PGS-EA | | | | PGS-GI | | | |
|---|---|---|---|---|---|---|---|---|---|---|---|---|---|---|---|---|---|---|---|---|---|---|---|
| **UV** | **a** | **b** | **ab** | **UV** | **a** | **b** | **ab** | **UV** | **a** | **b** | **ab** | **UV** | **a** | **b** | **ab** | **UV** | **a** | **b** | **ab** | **UV** | **a** | **b** | **ab** |
| SA_4 | 0.094 | 0.063 | 0.006 | SA_MIP | 0.083 | 0.108 | 0.009 | CT_FOP1 | 0.115 | 0.122 | 0.014 | | | | | DWI_LO1 | 0.136 | -0.050 | -0.007 | DWI_45 | 0.054 | 0.305 | 0.017 |
| SA_MIP | 0.076 | 0.040 | 0.003 | SA_PH | 0.084 | 0.110 | 0.009 | CT_PreS | 0.069 | 0.105 | 0.007 | | | | | DWI_LO2 | 0.108 | -0.102 | -0.011 | DWI_Pir | 0.054 | -0.203 | -0.011 |
| SA_1 | 0.059 | -0.226 | -0.013 | SA_IP1 | 0.084 | 0.035 | 0.003 | CT_25 | 0.075 | -0.123 | -0.009 | | | | | DWI_5mv | 0.125 | -0.251 | -0.031 | | | | |
| SA_6r | 0.067 | 0.047 | 0.003 | | | | | | | | | | | | | DWI_6ma | 0.140 | 0.153 | 0.021 | | | | |
| SA_IFSa | -0.073 | 0.119 | -0.009 | | | | | | | | | | | | | DWI_44 | 0.136 | 0.068 | 0.009 | | | | |
| SA_OFC | 0.091 | 0.060 | 0.005 | | | | | | | | | | | | | DWI_45 | 0.111 | 0.164 | 0.018 | | | | |
| SA_OP1 | 0.070 | 0.055 | 0.004 | | | | | | | | | | | | | DWI_6r | 0.124 | 0.136 | 0.017 | | | | |
| SA_STGa | 0.042 | 0.073 | 0.003 | | | | | | | | | | | | | DWI_47s | 0.139 | 0.234 | 0.033 | | | | |
| SA_PH | 0.099 | 0.058 | 0.006 | | | | | | | | | | | | | DWI_43 | 0.132 | 0.159 | 0.021 | | | | |
| SA_IP1 | 0.070 | 0.066 | 0.005 | | | | | | | | | | | | | DWI_Pir | 0.092 | -0.210 | -0.019 | | | | |
| | | | | | | | | | | | | | | | | DWI_VMV1 | 0.125 | -0.157 | -0.020 | | | | |
| | | | | | | | | | | | | | | | | DWI_pOFC | 0.098 | -0.057 | -0.006 | | | | |
